# Zonal endothelial cell heterogeneity underlies murine renal vascular development

**DOI:** 10.1101/2025.01.13.632837

**Authors:** Peter M Luo, Neha Ahuja, Christopher Chaney, Danielle Pi, Aleksandra Cwiek, Zaneta Markowska, Chitkale Hiremath, Denise Marciano, Karen K Hirschi, M Luisa Iruela-Arispe, Thomas J Carroll, Ondine Cleaver

## Abstract

The renal vasculature consists of highly specialized blood vessels that carry out distinct physiological functions. Defining their transcriptional signatures and tracing their developmental ontogeny has thus far been challenging due to a lack of regionally specific endothelial biomarkers. Here, we performed single nuclear RNA sequencing (snucRNA-Seq) to interrogate embryonic renal endothelial cells. We identified nine different endothelial cell types, and validated expression of novel candidates using multiplex RNAScope. We interrogate biological characteristics, including cell cycle stage, of developmental blood vessels with a variety of *in* vivo tools. Using genetic tools based on expression of tip cell marker *Esm1*, we performed detailed lineage tracing, demonstrating multi-clonal and multi-directional endothelial contribution to glomerular vasculature. Together, this study provides the first validated, tool-focused developmental atlas of the murine renal vasculature and elucidates several novel cellular mechanisms of nephron vascularization.

## INTRODUCTION

The renal vasculature consists of several specialized vessel types which are required for proper kidney function.^1^ These vessels include the glomerular capillaries, vasa recta, and peritubular capillaries, which respectively filter, concentrate, and reabsorb and secrete solutes into the urine.^2^ Despite the recognition that organotypic and regionalized vascular beds likely profoundly alter therapeutic considerations, the developmental mechanisms underlying this heterogeneity, particularly in the kidney, remain understudied.^3^

During kidney development, stromal, vascular, and epithelial components all grow concomitantly. To date, study of kidney development has primarily focused on epithelial lineages involved in nephron formation.^4^ Briefly, reiterative reciprocal induction events between epithelial and mesenchymal components direct their contribution to different parts of the nephron and collecting duct system. These cellular progenitors are located at the kidney periphery, termed the *nephrogenic zone* (NZ). Kidney development progresses centripetally, with newer nephrons added superficially to older nephrons that extend into the kidney as they differentiate.^5^ These growing tissues likely form local microenvironments that direct regionally tailored blood vessel formation.

While distinct mechanisms of vascular growth have been described in other systems,^6,7^ mechanisms of kidney vascular development remain largely unclear. We and others have characterized morphological features of the developing vasculature in relation to nephrogenesis, including differential vascular density between the NZ and underlying cortex.^8–10^ While many models of kidney vascular development have been proposed, the cellular mechanisms by which developmental vascular plexuses give rise to mature vessel types, such as glomerular or peritubular capillaries, remains largely unclear.

Notably, glomerular vascularization, while of acute interest for *ex vivo* organogenesis,^11^ remains poorly understood due to the absence of useful markers and genetic tools. Glomerular endothelial cells (ECs) were first identified on 2D sections in ‘S-shaped bodies’, the penultimate stage of nephrogenesis before glomerular formation, in a ‘glomerular cleft’ adjacent to immature podocytes.^12^ Thus, the prevailing dogma has been that an angiogenic sprout invading the glomerular cleft represented the first vascularization event in nephrogenesis. However, we recently characterized ECs associated with early stages of nephrogenesis using 3D imaging, showing that the S-shaped body was enwrapped in a dense vascular net, rather than invaded by a single vessel.^8^ The specific cellular events coordinating nephron development and specialization of vessels during nephron vascularization remain of great interest.

Recent efforts have examined renal EC heterogeneity through single cell sequencing.^9,13,14^ However, within whole-organ transcriptomic datasets, ECs make up a relatively small population, making it difficult to identify unique subtypes.^15^ Multiple recent studies have sequenced antibody-enriched renal ECs, generating lists of marker genes for both embryonic^9^ and adult kidney vascular beds.^13^ However, the specificity of many proposed vessel markers remains unvalidated *in vivo*. In particular, molecular markers of the specific vessel beds unique to the developing kidney vasculature are lacking. Moreover, lineage relationships between developmental and mature vascular beds have not been elucidated *in vivo* in the kidney, as they have in other organs such as the heart.^16^

Here, we generate a developmental-specific transcriptional atlas of E18.5 kidney ECs. Importantly, we utilize a genetically-encoded fluorescent sorting strategy on isolated nuclei, which avoids cell loss during antibody/bead labeling techniques and prevents contamination of mural cell-endothelial doublets.^15^ Using multiplex RNAscope, we extensively validate regionalized, heterogeneous endothelial gene expression and identify markers of transient, development-specific vessel types. We further describe angiogenic events within the kidney and perform detailed lineage tracing to support a multi-clonal model of glomerular vascularization by these cells. In sum, this work presents a developmental atlas of the murine renal vasculature and elucidates several new cellular mechanisms of nephron vascularization.

## RESULTS

### The embryonic kidney vasculature comprises several specialized vessel types

To assess basic kidney vascular architecture at E18.5, we performed whole mount immunofluorescence (WMIF) for VE-cadherin (VECAD), a pan-EC marker, and Endomucin (EMCN), a capillary/venous marker (**Fig S1A**). Arteries notably lacked EMCN (**Fig. S1A**, white arrowhead) and were flanked by EMCN+ veins, as previously described.^8^ The afferent arterioles and descending vasa recta (DVR), which exhibit a more arterial phenotype than the efferent arteriole or ascending vasa recta (AVR), were also EMCN-negative (**Fig. S1A’-S1A”**, red/yellow arrowheads).^17,18^

Three zones of cortical capillary vasculature were clearly identified (**Fig. 1A**). Peritubular capillaries (PTCs) surrounded LTL+ proximal tubules in deepest parts of the cortex (**Fig. S1A”’**). Peripheral capillaries within the *nephrogenic zone* (NZ) were relatively sparse, especially compared to immediately underlying capillaries (**Fig S1A’’’’,** dotted line), as previously characterized at E13.5.^8,9^ Here, we term the sparser, peripheral plexus the *nephrogenic zone plexus* (NZP), and the denser capillaries network between the NZP and PTCs as the *sub-nephrogenic plexus* (SNZP).

**Figure 1.**
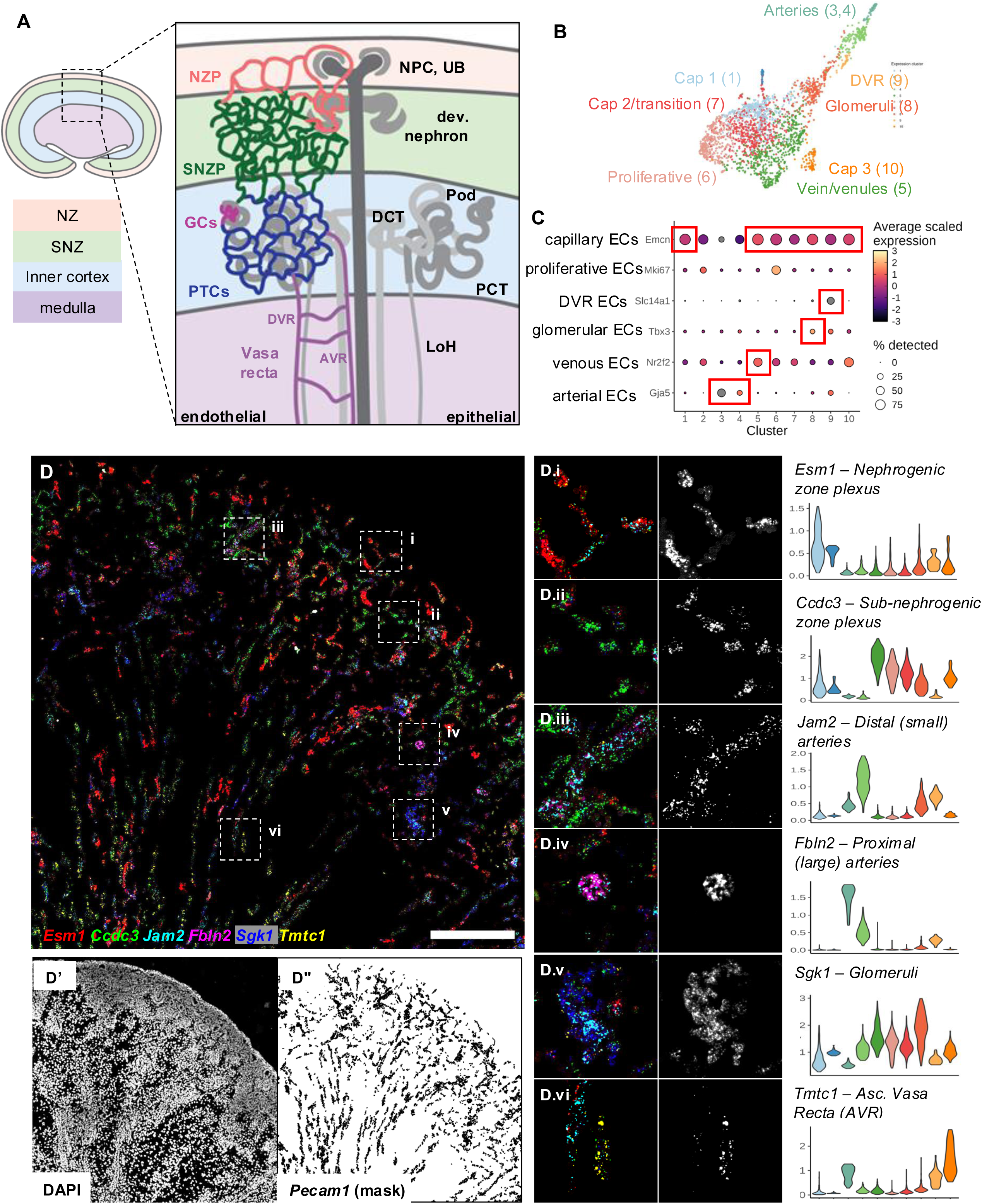
Single nuclear sequencing of E18.5 murine renal endothelial cells reveals heterogeneity and zonal patterns of expression. (A) Schematics of zonation of developing kidney (left) and regionalized vessel beds and associated epithelia (right). NZ: nephrogenic zone. SNZ: sub-nephrogenic zone. NZP: nephrogenic zone plexus. SNZP: sub-nephrogenic zone plexus. GC: glomerular capillary. PTC: peritubular capillary. AVR, DVR: ascending, descending vasa recta. NPC: nephron progenitor cell. UB: ureteric bud. Pod: podocyte. PCT, DCT: proximal, distal convoluted tubule. LoH: loop of Henle. (B) Dot plot of expression of known marker genes of vessel subtypes. (C) UMAP of endothelial nuclei with identified clusters annotated. (D) Hiplex RNAscope of 6 novel marker genes (*Tmtc1, Ccdc3, Esm1, Sgk1, Fbln2, Jam2*) at E18.5, masked using *Pecam1* RNAscope. (D’) DAPI channel, showing bounds of kidney section. (D”) 5um- diameter *Pecam1* mask, created using IMARIS. (D.i-D.vi) Zoom of (D), showing various vessel types. (D.i’-D.vi’) Single channel image of select marker gene. (D.i”-D.vi”) Violin plot of relative expression of marker genes within all clusters. Scale bars: 200µm (D).

### Single nuclear sequencing of E18.5 renal ECs reveals known and novel heterogeneity

To develop a transcriptional atlas of the developing mouse kidney vasculature, we utilized a *Flk1::H2B-eYFP* reporter to label EC nuclei.^19^ The reporter was active in most endothelial (ERG+) nuclei at E18.5 (**Fig. S1B-C**), allowing isolation of renal ECs following dissection of embryonic kidneys. YFP immunostaining was lower, however, in COUP-TFII+ venous ECs (**Fig. S1C-F**).^20^ We isolated and enriched endothelial nuclei from E18.5 kidneys using fluorescence assisted cell sorting (FACS), and we sequenced 2,447 nuclear transcriptomes using a 10x Genomics platform. Clustering revealed 9 subtypes of endothelial nuclei (after discounting cluster 2 due to lack of EC-specific markers) (**Fig. 1B**). We used enrichment of previously reported markers *Gja5, Nr2f2*, *Mki67*, *Tbx3*, and *Slc14a1* to identify arterial (cluster 3,4), venous (cluster 5), proliferative (cluster 6), glomerular (cluster 8), and DVR (cluster 9) nuclei, respectively (**Fig. 1C, Data S1**).^9,20–23^ In addition, three clusters of capillary nuclei exhibited high *Emcn* expression (**Fig. 1B-C**, cluster 1, 7, 10). Differentially expressed marker genes for each cluster were identified (**Table S1**).

To validate marker gene expression *in vivo*, we performed multiplex RNAscope fluorescent *in situ* hybridization at E18.5 (**Fig. 1D**). Non-endothelial signal was excluded using a 5μm mask around *Pecam1* probe signal (**Fig. 1D”)**. Significant non-endothelial expression of marker genes, as well as full gene names and reported roles in ECs, are summarized in **Table S2**. We identified distinct vessel beds with enriched expression of each marker gene, validating new markers of known vessel types and identifying remaining clusters based on regionalized marker gene expression (**Fig. 1D.i-1D.vi**).

We characterized the heterogeneity of various kidney vessel types using RNAscope. We performed IF for PODXL (marking podocytes) to identify glomeruli (**Fig. 2A**),^24^ along with RNAscope for cluster 8-enriched glomerular markers (see **Data S1** for UMAPs) (**Fig. 2B)**. *Sgk1* was enriched in glomerular ECs within the cortical vasculature (**Fig. 2B, B’**). Notably, *Sgk1* expression was lower in nascent glomeruli, which were more superficial and exhibited cuboidal podocytes (**Fig. 2C-D’**). *Chrm3* was expressed specifically in ECs of both immature and mature glomeruli (**Fig. 2B”**), but was not homogeneously expressed within glomeruli (**Fig. 2C”-D”**). Notably, *Esm1,* while not expressed exclusively in cluster 8 (**Data S1**), was expressed in a subset of *Chrm3-* negative glomerular ECs (**Fig. 2B”’-D’”**).

**Figure 2.**
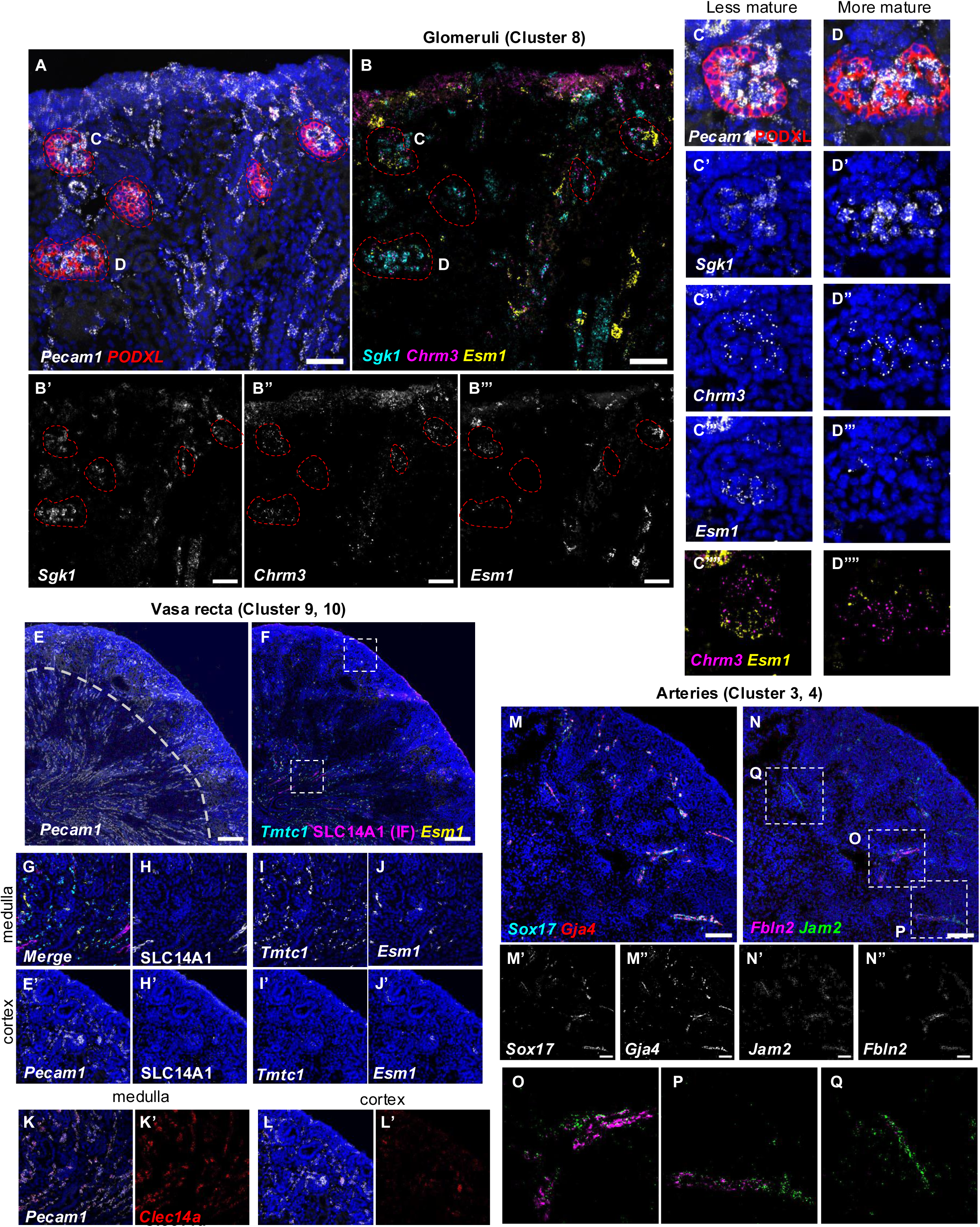
Novel markers and heterogeneity found within renal glomerular, vasa recta, and arterial vasculature. (A) *Pecam1* RNAscope and PODXL IF of E18.5 control kidney cortex. Red dotted lines, glomeruli. (B) *Sgk1, Chrm3, Esm1* RNAscope on section from (A). (B’-B”’) Single channel images. Arrows cortical vessels. (C-D””) Zoom of (A), showing single glomeruli. (C’-D”’’) Single channel zooms of B. (E) *Pecam1* RNAscope of E18.5 kidney. Dotted line: corticomedullary junction. (F) *Tmtc1* (masked) and *Esm1* RNAscope with SLC14A1 IF on section from (E). (G) Medullary zoom of (F). (H-J) Single channel views of (G). (E’, H’-J’) Cortical zoom of (E) and single channel views of (F). (K) *Pecam1* and *Clec14a* RNAscope of E18.5 medulla, as in (G). (K’) Single channel view. (L) *Pecam1/Clec14a* RNAscope of E18.5 cortex, as in (E’). (L’) Single channel view. (M) *Sox17* and *Gja4* RNAscope of E18.5 kidney cortex. (M’-M”) Single channel views. (N) *Fbln2* and *Jam2* RNAscope of E18.5 section from (M). (N’-N”) Single channel views. (O-Q) Zoom views of (N), showing individual arteries. Scale bars: 50µm (A-B), 200µm (E-F), 100µm (M-N).

Vasa recta capillaries were located within the medulla of E18.5 kidneys (**Fig. 2E**, within dotted line). DVR capillaries expressed SLC14A1 by IF as previously characterized,^23,25^ which was enriched in cluster 9 (**Fig. 2F, H, Data S1**). *Tmtc1*, which was enriched in cluster 10, was expressed specifically in a subset of medullary ECs with minimal cortical endothelial expression (**Fig. 2F, I-I’, Data S1**). *Tmtc1* and SLC14A1+ vessels did not overlap, suggesting that *Tmtc1*+ vessels are AVR capillaries, for which current markers are lacking.^9^ Notably, a subset of *Tmtc1-*negative, SLC14A1-negative medullary ECs expressed *Esm1*, our cluster 1 marker which was also expressed at the kidney periphery (**Fig. 2F, J-J’**). Interestingly, *Clec14a*, which was enriched in both clusters 9 and 10, was expressed in all vasa recta ECs, with low cortical endothelial expression (**Fig. 2K-L, Data S1**).

Pan-arterial markers *Sox17* and *Gja4* were expressed in clusters 3 and 4 (**Fig. 2M-M”, Data S1**).^26,27^ *Fbln2* (cluster 3) and *Jam2* (cluster 4) were differentially expressed in proximal (larger) arteries and distal arteries/arterioles, respectively (**Fig. 2N-Q, Data S1**). Notably, some arteries transitioned from primarily expressing *Fbln2* to expressing *Jam2* as they traversed to more cortical regions, indicating a transcriptional gradient along the arterial tree (**Fig. 2P**). Taken together, these results validate novel marker genes of glomerular, vasa recta, and arterial ECs, and reveal previously uncharacterized heterogeneity within these developing vessel beds.

The marker genes of two remaining clusters, *Esm1* (cluster 1) and *Ccdc3* (cluster 5) displayed interesting, restricted, expression patterns near the edge of the cortex (**Fig. 1D**). In addition to glomerular and medullary expression, *Esm1* was expressed in the peripheral NZP (**Fig. 1D.i**). Despite cluster 5 being enriched for venous genes such as *Nr2f2*, *Ccdc3* expression was enriched in the SNZP immediately deep to the NZ (**Fig. 1D.ii**). These results demonstrate distinct zonal heterogeneity and validate specific molecular markers for the developing kidney vasculature.

### Embryonic endothelial heterogeneity partially perdures in post-nephrogenic kidneys

To test whether these molecular markers remain regionally restricted post-natally,^9^ we analyzed their expression at P16. Consistent with higher expression in older glomeruli at E18.5, *Sgk1* was highly enriched in glomeruli (**Fig. S2A-B, B”**). *Fbln2* and *Jam2* remained differentially enriched in proximal and distal arteries, respectively (**Fig. S2A-B”’**). Notably, *Fbln2* was also expressed in large veins, suggesting that its expression correlates with large vessel maturity independent of arteriovenous identity (**Fig. S2D,** arrowhead). *Tmtc1* remained restricted to the medullary AVR (as opposed to *Jam2+* DVR) (**Fig. S2A, C**). No markers that we identified had specific expression within peritubular capillaries. Notably, expression of *Ccdc3* and *Esm1* change drastically between E18.5 and P16. At P16, *Ccdc3* expression was lower with minimal endothelial-specific expression (**Fig. S2E**), while *Esm1* was restricted to the outer medulla (**Fig. S2F**), This contrasts with the cortical plexuses that display restricted *Ccdc3* and *Esm1* expression at E18.5.

### Stromal loss of netrin 1 results in patterning defects of the sub-nephrogenic and nephrogenic zone vessels

Having established a molecular map of developing kidney ECs, we aimed to apply it to a model of altered renal vascular development. We and others recently reported vascular and mural cell defects resulting from netrin 1 (Ntn1) ablation from stromal progenitors in the NZ (*Foxd1*^GFPCre^; *Ntn1*^flox/flox^, abbreviated *Ntn1*^SPKO^).^28,29^ Specifically, we reported dense ectopic vasculature at the surface of E13.5 *Ntn1*^SPKO^ kidneys, as opposed to the sparse NZP in controls. However, the identity of these vessels is unclear.

WMIF for EMCN on E15.5 *Ntn1*^SPKO^ kidneys revealed ectopic blood vessels along the kidney surface, similar to E13.5 (**Fig. S3A-B**). We previously reported arterial mispatterning at E13.5 within *Ntn1*^SPKO^ kidneys, however, ectopic blood vessels were broadly non-arterial by CX40 WMIF (**Fig. S3A-B”**). To assess transcriptional signatures of developing vascular plexuses within ectopic vessels, we performed RNAscope *for Pecam1, Esm1*, and *Ccdc3* (**Fig. 3A-D**). Notably, while *Ccdc3* and *Esm1* were both expressed in some, but not all ECs near the surface of mutant kidneys (**Fig. 3D.i-D.iii**), significantly more ECs in mutant kidneys lacked expression of either gene (**Fig. 3D-E**). Taken together, ectopic vessels in *Ntn1*^SPKO^ are highly heterogeneous, displaying neither distinct NZP nor SNZP character, indicating dysregulation of both vessel beds.

**Figure 3.**
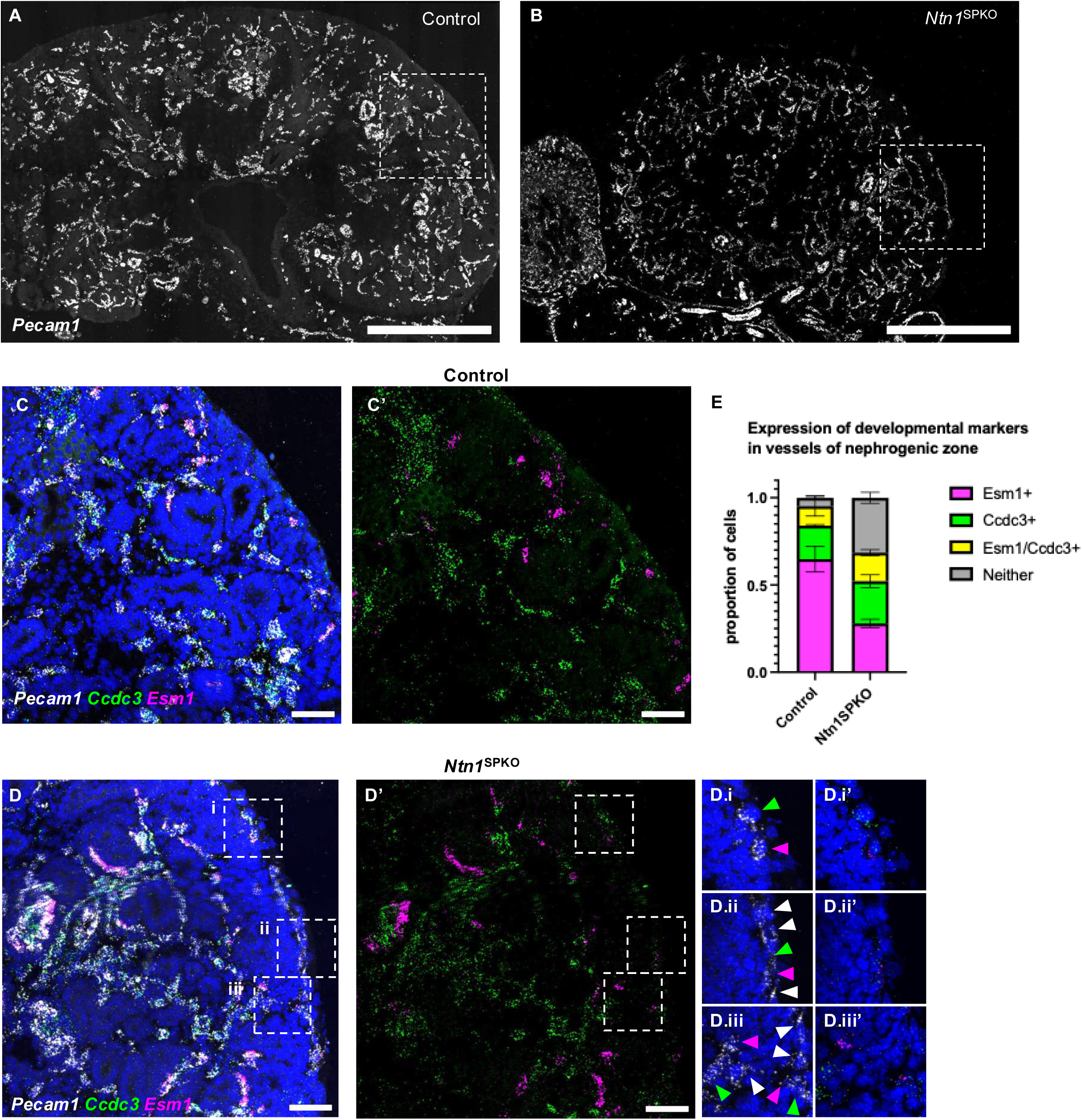
Loss of stromal netrin 1 results in altered developmental vascular organization at E15.5 as revealed by novel marker genes. (A-B) *Pecam1* RNAscope of E15.5 control (A) and *Ntn1*^SPKO^ (B) kidney sections. (C) Zoom of kidney periphery of control and (D) mutant kidney, with *Esm1* and *Ccdc3* RNAscope. (C’-D’) *Ccdc3* and *Esm1* RNAscope only. (D.i – D.iii) Insets of (D), showing individual vessels at the kidney periphery. Arrowheads: *Esm1*+ (magenta), *Ccdc3+* (green), or neither (white). (D.i’ – D.iii’) 2 channel view of D’, with dapi. (E) Quantification of proportion of nephrogenic zone (NZ) ECs classified by *Esm1/Ccdc3* positivity. n=155 nuclei, control (2 embryos); 418 nuclei, mutant (3 embryos). Error bars: SEM. Scale bars: 500µm (A-B), 50µm (C-D).

To assess potential downstream signaling, we analyzed expression of known Ntn1 receptors *Unc5b, Unc5c,* and *Neo1* in our transcriptomic dataset*. Unc5b* was enriched in NZP and arterial clusters,^30^ *Unc5c* was enriched in the SNZP cluster, and Neo1 was enriched in glomerular/arterial clusters (**Fig. S3C-E**). Surprisingly, no known receptors were expressed within ectopic vessels (**Fig. S3F-G**), suggesting that ectopic vessel growth in *Ntn1*^SPKO^ kidneys is not mediated by known cell-intrinsic Ntn1 receptor expression. Together, these results demonstrate the utility of our atlas in analyzing models of altered renal vascular development.

### Ccdc3 specifically marks a transient sub-nephrogenic plexus

Next, we characterized the dynamics and behavior of development-specific renal vascular beds. We performed WMIF for PAX2 (marking developing nephrons)^31^ and LTL at E15.5, E18.5, and P2 to define the boundaries of the sub-nephrogenic plexus throughout development (**Fig 4A-C**). The sub-nephrogenic plexus is restricted to development, as the distance between PAX2 and LTL staining diminishes rapidly near the end of nephrogenesis (**Fig. 4C**, white bracket).

**Figure 4.**
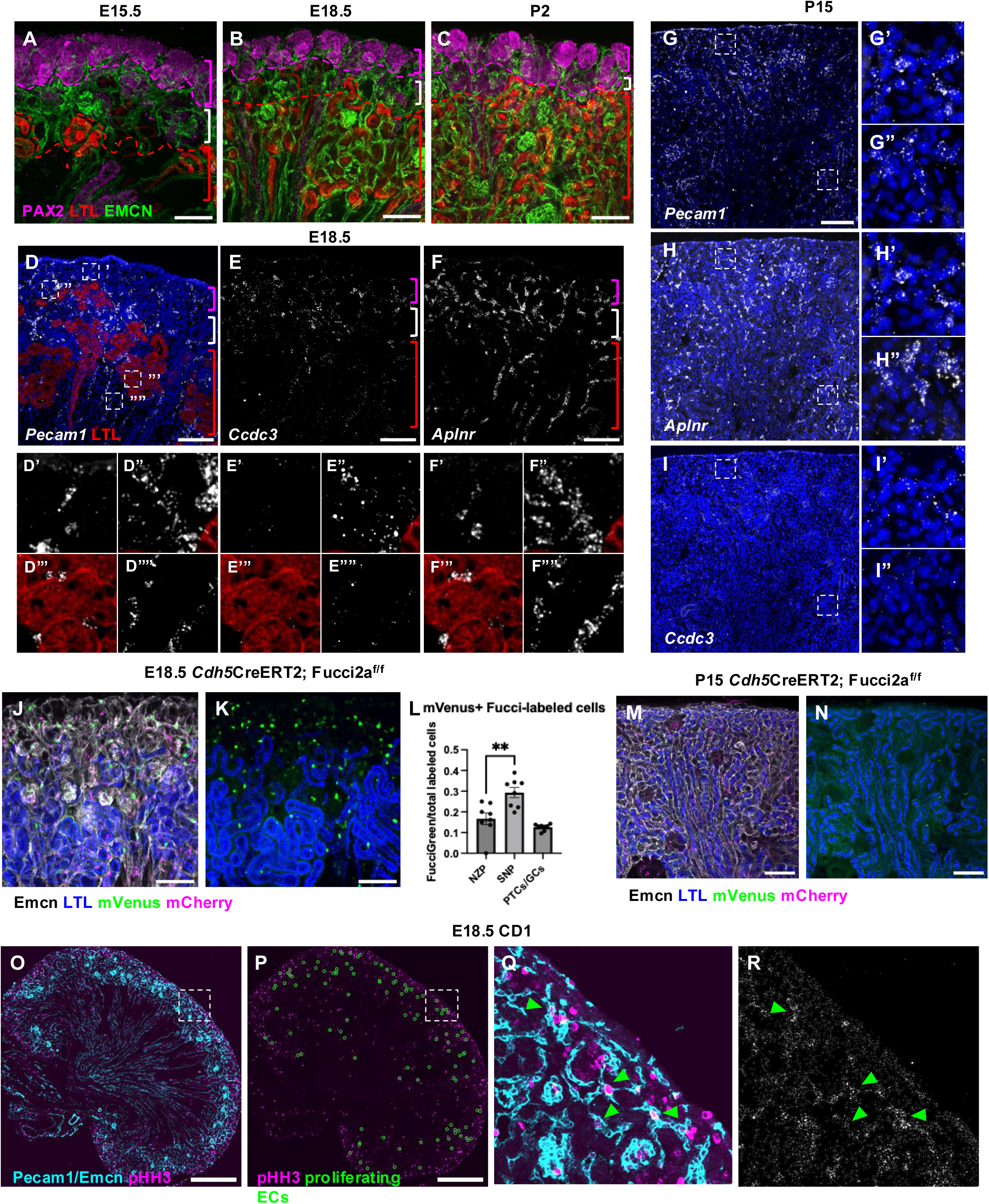
*Ccdc3* expression is restricted to the sub-nephrogenic plexus, a transient structure present during development. (A-C) Whole mount immunofluorescence (WMIF) of E15.5-P2 kidney sections for PAX2, LTL, and EMCN. Dotted lines: border of LTL+ proximal tubules (red) and PAX2+ nephrogenic epithelium (magenta). Brackets: magenta, NZ; white, sub-nephrogenic zone; red, medulla. (D) *Pecam1* RNAscope with LTL IF of cortical E18.5 section. (D’-D””) Zooms of NZ (‘), sub-nephrogenic plexus, SNZP (“), peritubular capillary, PTC (“’), and vasa recta vessels (“”), respectively. (E-F) *Ccdc3* and *Aplnr* RNAscope of E18.5 kidney cortex. (E’-E””, F’-F””) Insets, as in (D). (G-I) P15 kidney RNAscope of *Pecam1*, *Aplnr*, and *Ccdc3,* respectively. (G’-I’) Insets of outer cortex of panels shown in G-I. (G’’-I”) Insets of inner cortex of panels shown in G-I. (J) WMIF of E18.5 *Cdh5*CreERT2; Fucci2a cortical kidney section for LTL, EMCN, mVenus, and mCherry. (K) mVenus and LTL only. (L) Quantification of proportion of mVenus cells/total labeled cells in cortical vessel beds. Error bars: SEM. n=2,242 nuclei, 3 embryos. p=0.0041, unpaired t test. (M) WMIF of P15 *Cdh5*CreERT2; Fucci2a cortical kidney section for LTL, EMCN, mVenus and mCherry. (N) mVenus and LTL only. (O) IF of E18.5 CD1 kidney for PECAM1/EMCN and pHH3. (P) pHH3 only. Green circles: proliferating ECs identified visually with pHH3 and PECAM1/EMCN co-staining. (Q) Zoom of NZ. Green arrowheads: Proliferating ECs. (R) *Ccdc3* RNAscope of inset from (Q). Green arrowheads: proliferating ECs. Scale bars: 100µm (A-N), 500µm (O-P).

*Ccdc3,* our marker gene of cluster 5, was enriched in sub-nephrogenic plexus ECs, visualized using co-IF for LTL at E18.5 (**Fig. 4D-E**). *Ccdc3* was not expressed in NZ vessels, glomeruli, or arteries, and was minimally expressed in peritubular capillaries and vasa recta (**Fig. 4E’-E””**). Barry *et al.* proposed the presence of *Aplnr-*enriched renal vascular progenitors and validated APLNR protein in the sub-nephrogenic plexus by IF at E17.5.^9^ By RNAscope, *Aplnr* was less specific to the sub-nephrogenic plexus than *Ccdc3*, displaying appreciable expression in most vessels at E18.5, including peritubular capillaries and vasa recta (**Fig. 4F-F””**). Notably, at P15, after nephrogenesis concludes, cortical ECs exhibit strong, uniform *Aplnr* expression and significantly less *Ccdc3* expression (**Fig. 4G-I**).

### The sub-nephrogenic plexus is a vein-like niche of proliferating endothelial cells

Cluster 5 also displayed restricted expression of classical venous markers (**Data S1**). Large veins (adjacent to *Gja4+* arteries nearest the hilum) expressed very little *Ccdc3,* despite being topologically within the cortex (**Fig. S4A-A.i’**). Medium-sized veins along the corticomedullary junction contained *Ccdc3* expressing nuclei (**Fig. S4A.ii**). However, all lumenized veins expressed less *Ccdc3* than the sub-nephrogenic plexus, which was often associated with a *Gja4*+ arteriole (**Fig. S4A.iii**). WMIF at E18.5 for VECAD and EMCN revealed direct connections between venous lumens (**Fig. S4B-B”’,** red arrowheads) and the sub-nephrogenic plexus through serial z-planes (**Fig S4B-B”’,** yellow arrowheads), without connections to the nearby arteries. These observations suggest that the sub-nephrogenic plexus represents a distinct venous/venule identity within the developing kidney.

*Ccdc3* was also highly expressed in cluster 6 (proliferative ECs) (**Fig. 1C**).^32^ To visualize the zonation of proliferating ECs within the cortex, we utilized a *Cdh5*CreERT2-driven Fucci2a (*Cdh5*Fucci) reporter, which expresses mCherry-Cdt1 or mVenus-Geminin during G1 and G2/M phase, respectively (**Data S1**).^33–36^ At E18.5, the sub-nephrogenic plexus contained significantly more proliferative cells (G2/M phase, mVenus+) than either the NZP or LTL-associated PTC (**Fig 4J-L**). At P15, proliferative cells were very sparse in the cortex, indicating that the cortical endothelium is no longer highly proliferative (**Fig. 4M-N)**.

To validate EC proliferation within the sub-nephrogenic plexus and in relation to other cell types, we performed IF for pHH3, a G2/M phase marker, along with PECAM1/EMCN (**Fig. 4O**) and RNAscope for *Ccdc3*. While overall most pHH3+ cells were found in the NZ, pHH3+ ECs were enriched within the sub-nephrogenic plexus and expressed high *Ccdc3* (**Fig. 4P-R**, arrowheads). Together, these results demonstrate that the *Ccdc3-*enriched sub-nephrogenic plexus represents a proliferative venous structure that may support vascular growth during kidney development.

We also analyzed proliferation and cell cycle stage in other vessels. In both *Cdh5*Fucci and pHH3 IF, other commonly proliferative cells at E18.5 included glomerular endothelium and both AVR and AQP1+ DVR capillaries (**Fig. S4C-E**).^25^ At P15, mVenus+ proliferative cells are highly enriched in the outer medulla of *Cdh5*Fucci kidneys, in both DVR and AVR capillaries (**Fig. S4F-H**).

In the mouse retina, endothelial late G1 phase arrest marked by FucciRed/mCherry enrichment has been reported to drive arterial identity.^35^ We performed WMIF for mCherry and aSMA (to identify arteries) in E18.5 and P15 *Cdh5*Fucci kidneys, and observed that larger caliber proximal arteries, but not distal arteries and arterioles, were enriched for mCherry+ cells (**Fig. S4I-K**). Notably, at both E18.5 and P15, the medulla contained the most mCherry+ endothelial nuclei (**Fig. S4L, N)**. Both SLC14A1+ DVR and EMCN+ AVR capillaries displayed mCherry+ staining (**Fig. S8M, O**), indicating cell cycle control independent of arteriovenous identity. Together, these results demonstrate novel findings regarding regionalized cell cycle control within the kidney endothelium.

### Esm1 is expressed in the nephrogenic zone vasculature

Next, we aimed to characterize the development of the NZP, marked by *Esm1* enrichment in cluster 1. *Esm1+* NZ vessels displayed variable *Ccdc3* expression, but consistently expressed less *Ccdc3* than ECs in the SNZP (**Fig. 5A-B””**, white dotted line). To visualize the spatial relationship between these two zones of the cortical kidney vasculature, we analyzed superficial sections 10-30 μM from the edge of the kidney (a.k.a ‘edge section’) (**Fig. 5C-D**), such that the entire section would span the NZ. Most NZ vessels expressed *Esm1,* while fewer vessels expressed *Ccdc3* (**Fig. 5D**). However, multiple NZ vessels contained contiguous *Esm1+* and *Ccdc3+* nuclei, or *Esm1/Ccdc3* dual positive nuclei (**Fig. 5D’-D”’, S5A,** arrowheads). Notably, vessels found superficial to SIX2+ caps were only *Esm1+* (**Fig. 5D”’, S5A,** magenta arrow). Together, these results suggest that the *Ccdc3+* SNZP and the *Esm1+* NZP are bridged by ECs that variably co-express *Ccdc3* and *Esm1*.

**Figure 5.**
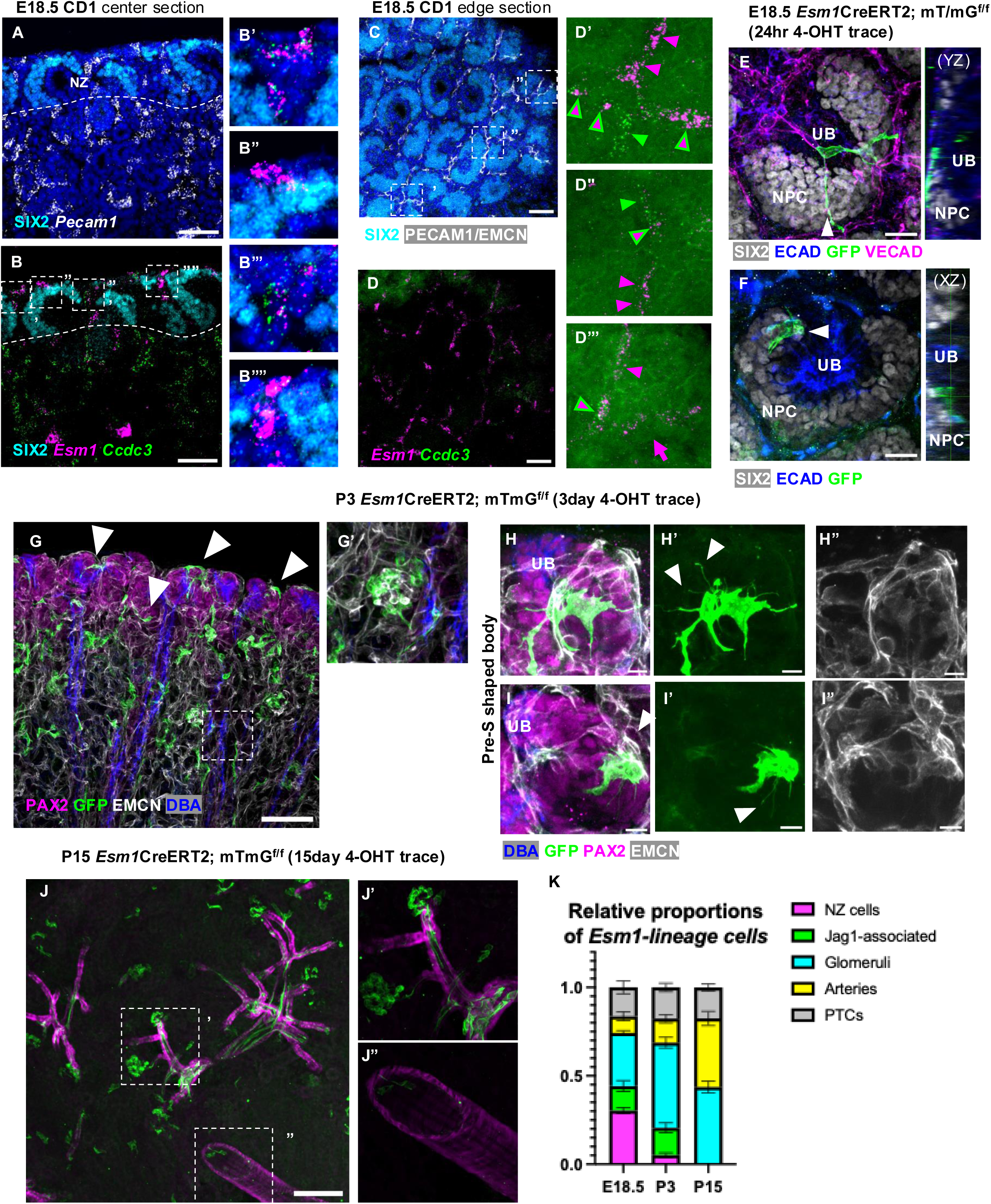
*Esm1* is expressed in nephrogenic zone plexus, whose endothelial cells contribute to arterial and glomerular vasculature. (A) *Pecam1* RNAscope with SIX2 IF of E18.5 CD1 kidney cortex. White dotted line: Border of NZ, containing SIX2+ NPCs. (B) *Ccdc3* and *Esm1* RNAscope with SIX2 IF on section from (A). (B’-B””) Zooms, showing individual NZ vessels. (C) PECAM1/EMCN and SIX2 IF on E18.5 edge section (within 30um of organ periphery). (D) *Ccdc3* and *Esm1* RNAscope on section from (C). (D’-D”’) Zooms, showing NZ vessels. Note that green channel is adjusted to ensure proper visualization of *Ccdc3* RNAscope signal. Arrowheads: *Esm1+* (magenta), *Ccdc3+* (green), *Esm1/Ccdc3+* (magenta with green border). Arrow: vessel spanning NPC cap. (E) WMIF of E18.5 *Esm1*CreERT2; mT/mG edge kidney section (24-hr pulse trace) for SIX2, VECAD, ECAD, and GFP. UB, NPC: ureteric bud, nephron progenitor cells. White arrowhead: GFP+ filopodia. Orthogonal view (YZ) of plane indicated by arrowhead. (F) WMIF of different area of same section as (E). Arrowhead: GFP+ tip cell. Orthogonal view (XZ) of plane indicated by arrowhead. NPC, UB: nephron progenitor cell, ureteric bud. (G) WMIF of P3 *Esm1*CreERT2; mT/mG kidney (3-day lineage trace) cortical section for PAX2, GFP, EMCN, and DBA. Arrowheads: PAX2-associated GFP+ cells. (G’) Zoom of (G), showing glomerulus. (H-I) WMIF of P3 *Esm1*CreERT2; mT/mG (3-day lineage trace) kidney section for DBA, PAX2, EMCN and GFP, showing developing nephrons before the S-shaped body stage. Arrowheads: GFP+ filopodia. (H’-I”) GFP, EMCN only. (J) WMIF of P15 *Esm1*CreERT2; mT/mG (15-day lineage trace) cortical section for GFP and aSMA. (J’) Zoom of distal arterial tree and glomeruli. (J”) Zoom of proximal artery. (H) Quantification of proportions of *Esm1+* nuclei (by RNAscope at E18.5 CD1) or GFP+ cells (by IF at P3/P15 on lineage traced tissue) within different vessel types. Error bars represent SEM. n=568 nuclei, 2 embryos (E18.5), 765 cells, 2 embryos (P3), 497 cells, 2 embryos (P15). Scale bars: 50µm (A-D), 20µm (E-F), 100µm (G,J), 10µm (H-I).

### Esm1-labeled tip cells are found in the nephrogenic zone plexus

*Esm1* classically marks ‘tip’ cells, which are specialized sprouting cells that utilize filopodia to sense microenvironmental growth factors and direct vessel growth at angiogenic fronts.^37^ Other tip cell markers, such as *Cxcr4, Dll4,* and *Adm*,^38–40^ were enriched in cluster 1, suggesting that NZP contains angiogenic tip cells (**Data S1**).

To assess the location and morphology of putative *Esm1+* tip cells within the NZ plexus, we utilized a *Esm1*CreERT2; mT/mG mouse to induce membrane GFP at E17.5 with 4-hydroxytamoxifen (4-OHT), assessing at E18.5 for a 24-hour short pulse. Validating our system, GFP+ cells in E18.5 short-pulse *Esm1*-lineage traced kidneys were restricted to the same vessel beds that expressed *Esm1* by RNAscope **(Fig. S5A**). WMIF on edge sections revealed GFP+ cells with long filopodia in the NZ, suggesting that labeled cells at the organ periphery were indeed tip cells (**Fig. 5E-F**). NZ tip cells were found in multiple configurations in relation to other cellular compartments. Some GFP+ cells were superficial to the SIX2+ nephron progenitor cap cells (NPCs), sending projections that spanned the NPC cap, bridging vessels at the surface of the kidney (YZ orthogonal view, **Fig. 5E**). These tip cells fit previous descriptions of the polygonal morphology and cyclical development of the NZP.^10^ By contrast, other GFP+ tip cells localized deeper in the plexus, positioned between the NPC cap and the UB (**Fig. 5F**, XZ orthogonal view). These cells surrounded a nascent renal vesicle (unstained structure between NPC cap and UB) with filopodia (**Fig. S5B-C,** inset), demonstrating tip cell association with the earliest stages of nephrogenesis.

### The cortical Esm1 tip cell lineage contributes specifically to glomeruli and arteries

To assess the fate of *Esm1+* tip cells, we performed lineage tracing of NZ tip cells using the *Esm1*CreERT2; mT/mG mouse, induce sparse membrane GFP labeling at P0 using 4-OHT and assessing GFP localization at P3 and P15 (**Data S1**). At P3, GFP+ cells were found in glomeruli within the cortex (**Fig. 5G, G’**). Additionally, in more peripheral regions of the cortex, GFP+ cells associated tightly with PAX2+ developing nephron epithelium (**Fig. 5G**, arrowheads). Single GFP+ cells with highly complex filopodial morphology spanned PAX2+ early stage developing nephrons, indicating that *Esm1*-lineage traced cells may retain a tip-cell like morphology over time (**Fig. 5H-I**, arrowheads). GFP+ cells were found in multiple locations in relation to PAX2+ epithelium. Notably, by P15, cortical *Esm1*-lineage cells were primarily located in glomeruli or distal arteries (**Fig. 5J-J’**), but not proximal arteries (**Fig. 5J”**).

To track the progression of the cortical *Esm1-*lineage, we classified the *Esm1* lineage as either NZP, JAG1+ epithelium-associated, glomerular, arterial, or other (including peritubular capillaries) (**Fig. S5D-F**), using RNAscope at E18.5 and GFP+ *Esm1-*lineage cells at P3 and P15. JAG1 is a specific marker of the nephrogenic epithelium that is absent from NPCs.^41^ At E18.5, 30% of *Esm1*+ nuclei were in the NZP (**Fig. 5K**). Through development, *Esm1*-derived cells progressively diminished in NZ tip cells, then in JAG1-associated cells, while correspondingly increasing in glomerular and arterial cells (**Fig. 5K**). Importantly, the proportion of other GFP+ vessel labeling, including peritubular capillary cells, did not increase, suggesting that cortical tip cells in the NZ or associated with developing nephrons do not contribute to PTCs.

### Glomerular vasculature arises from multiple endothelial sources

To understand the cellular mechanism by which the *Esm1-*lineage participates in glomerular vascularization, we analyzed P3 lineage traced tissue at ‘comma-shaped body’ (CSB) or ‘S-shaped body’ (SSB) late stages of nephrogenesis. GFP+ cells were tightly associated with multiple ‘segments’ of the developing nephron. As classically expected, angiogenic *Esm1*-lineage cells were frequently found in the ‘glomerular cleft’, located between immature podocytes and medial segment of the CSB or SSB (**Fig. 6A-D**).^12,42^ In earlier stage SSBs, these cells extended filopodia into the cleft and towards the podocytes (**Fig. 6A”-B”, S6B-B’**). As SSBs progressed, GFP+ cells in the glomerular cleft no longer exhibited filopodia, but adopted a spoon-like morphology to envelop the medial SSB segment as part of a previously described vascular net (**Fig. 6C**).^8^ Importantly, GFP+ cells formed connections with other vessels under the podocyte lip of the SSB (**Fig. 6C’-E’**, arrowheads). As SSBs transitioned into immature glomeruli, we observed formation of a clear vascular tube within the glomerular cleft, with a recognizable inflow and outflow (**Fig. 6E’)**. This tube was only partially composed of GFP+ cells, and WMIF for SOX17 revealed that this vascular tube was arteriolar, suggesting that GFP+ cells in the glomerular cleft contribute to efferent or afferent arterioles (**Fig. 6E’-E”**).

**Figure 6.**
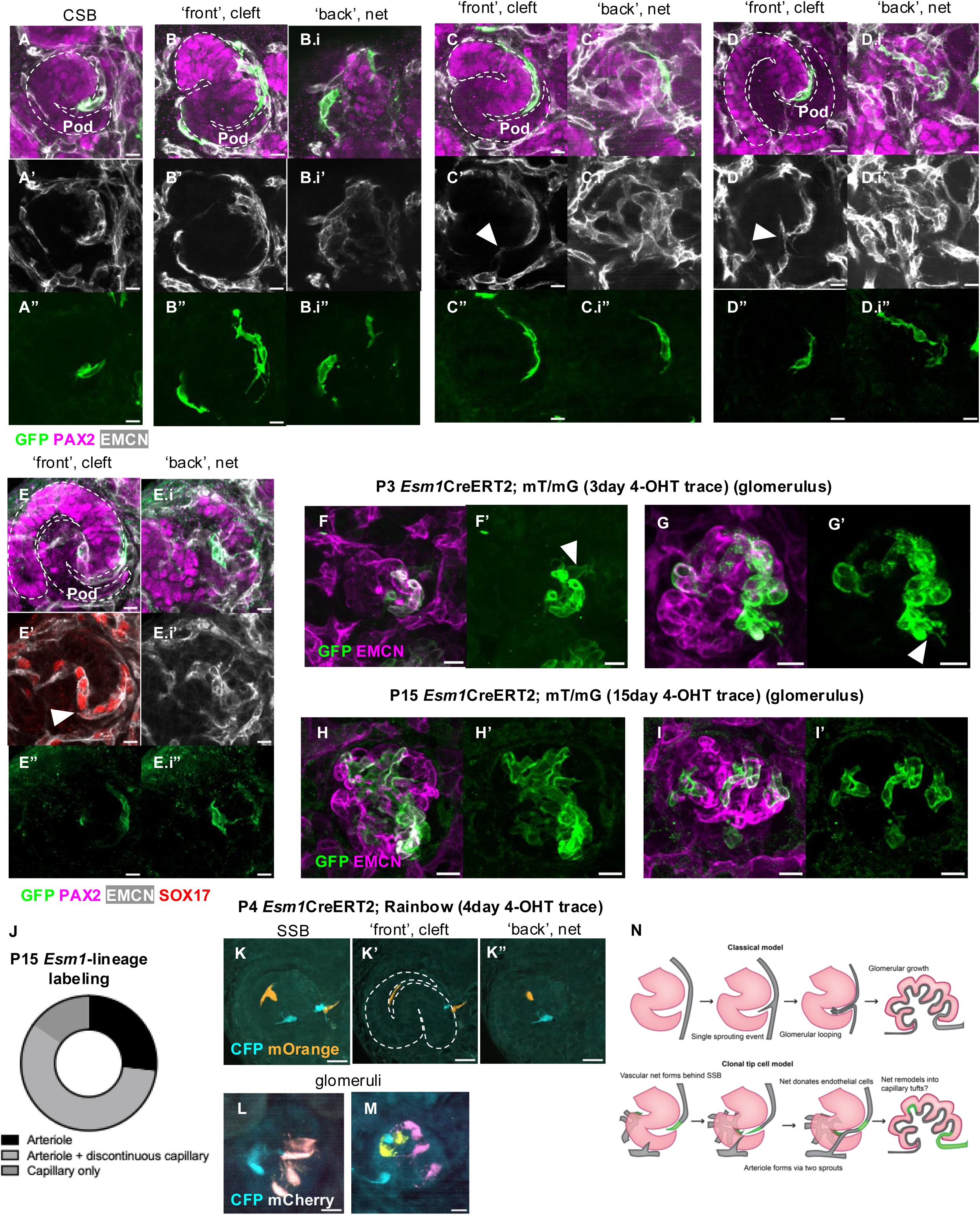
*Esm1+* tip cells in the nephrogenic zone vasculature contribute to glomerular vessels in a coordinated, multi-clonal manner. (A) WMIF of P3 *Esm1*CreERT2; mT/mG section, showing comma shaped body (CSB) for GFP, PAX2, and EMCN. (A’) EMCN only. (A”) GFP only. (B-E) WMIF as in (A), showing front half of S-shaped body (outlined) stages with the glomerular cleft. Pod: podocytes. Arrowheads: endothelial connections across podocytes, indicating arteriole formation. (B’-E’) EMCN only. (B”-E”) GFP only. (B.i-E.i) WMIF of back half of SSB in (B-E), showing vascular net. (B.i’-E.i’) EMCN only. (B.i”-E.i”) GFP only. (F-G) WMIF as in (A-E), showing nascent glomeruli. Arrowheads: GFP+ arteriole. (F’-G’) GFP only. (H-I) WMIF of P15 *Esm1*CreERT2; mT/mG sections, showing glomeruli. (H’-I’) GFP only. (J) Quantification of proportion of glomeruli at P15 with continuous, discontinuous, or isolated GFP+ cell labeling. n=25 glomeruli, 2 embryos. (K) 3D image of native fluorescence of P4 *Esm1*CreERT2; Rainbow reporter (induced with 4-OHT at P0), showing SSB. (K’) Single Z plane, showing glomerular cleft. (K”) Single Z plane, showing back of SSB. (L-M) 3D native fluorescence as in (K), showing glomeruli. (N) Model of tip cell vascularization of glomeruli. Scale bars: 10µm (A-I,L-M), 20µm (K).

GFP+ cells also formed tight associations with other portions of the SSB (**Fig. 6B.i, 6D.i-E.i**). Notably, GFP+ cells were observed extending filopodia ‘behind’ the medial segment of the SSB (**Fig. 6B.i”)**. ECs formed a dense plexus as the medial segment outpouches ‘forward’ to give the SSB its 3D structure, variably containing GFP+ cells (**Fig. 6C.i’-6E.i’, 6C.i”-6E.i”**). Intriguingly, GFP+ cells in this vascular plexus did not appear to contribute to arterioles, but were found in close proximity, suggesting that they may incorporate into glomerular vasculature (**Fig. 6D.i”-E.i”**).

Stromal cells, including mesangial cells (a specialized type of pericyte), play an important role in glomerular function.^43–45^ We assessed incorporation of NG2+ cells using *Cspg4-*mEGP reporter mice at SSB stages at E17.5. At SSB stages with distinct arterioles visible by PECAM1 WMIF, GFP+ cells were found associated with endothelial cells within the glomerular cleft, analogous to *Esm1*-lineage cells within the glomerular cleft and contributing to a portion of the arteriolar vasculature (**Fig. S6C**). In addition, *Cspg4*-mEGFP+ cells were found densely associated with the vascular net (**Fig. S6C.i**). Together, these results show the close association of NG2-expressing cells, representing mural, pericyte, or mesangial cell types of stromal origin, associated with endothelial cells early in glomerular development.

### Esm1-lineage tip cells contribute in a mosaic and multi-clonal fashion to glomerular vasculature

When assessing glomeruli deeper in the cortex of P3 lineage traced tissue by WMIF, GFP+ cells rarely homogeneously labeled the entirety of EMCN+ glomerular capillary vasculature (**Fig. 6F’-G’**, arrowheads). GFP+ labeling frequently included a SOX17+ arteriole (**Fig S6C**), supporting our findings that cells within the glomerular cleft of SSBs form a vascular tube that becomes arterioles. Intriguingly, some glomeruli contained multiple discontinuous clusters of GFP+ cells that were separated by non-GFP+ vasculature, and other glomeruli contained a single cell or cluster of labeled cells within glomerular tufts (**Fig. 6G**).

To exclude the possibility that discontinuous or isolated GFP labeling was due to later induction by residual 4-OHT at the glomerular stage, rather than incorporation of SSB-associated tip cells, we analyzed glomeruli in P15 lineage traced tissue. GFP labeling continued to be mosaic in the majority of glomeruli analyzed, including discontinuous GFP labeling or labeling of single cells within the glomerular endothelium (**Fig. 6H-J**). Together, these results suggest that the *Esm1*-lineage labels different parts of the glomerular vasculature that remain interspersed by non-GFP+ endothelium.

Throughout nephrogenesis, the *Esm1* tip cell lineage contains various cellular configurations around epithelial components as described above. Labeling in different locations around SSBs was stochastic and did not always occur in conjunction. In addition, discontinuous and mosaic labeling of glomeruli suggest a multi-clonal contribution of the *Esm1-*lineage. To test this possibility, we next utilized a *Esm1*CreERT2; Rainbow reporter,^46^ where tri-color recombination was induced with 4-OHT at P0 and assessed at P4 (**Data S1**). At SSB stages, cells within the cleft and in the ‘back’ of the SSB, presumably within the vascular net, were labeled with different fluorophores and arose from different clones (**Fig. 6K**). Glomeruli were observed containing multiple labeled clones, demonstrating multiple tip cell contributions (**Fig. 6L-M**).

### Cortical Esm1 lineage contributes to distal arteries

Beyond contribution to arteriolar and capillary vasculature of the glomerulus, we also found that GFP+ cells contributed to distal arteries at P15. WMIF for GFP and SOX17 on P3 lineage traced tissue revealed SOX17+ distal arteries near the kidney periphery that contained GFP+ *Esm1-*lineage cells (**Fig. S6E**), as well as nearby GFP+ cells with lower SOX17 expression with filopodial connections to the artery (**Fig. S6E’-E”,** arrowheads), suggesting that these peripheral, NZ ECs may become arterial.

### Esm1 lineage in the medulla contributes to descending vasa recta

At E18.5, *Esm1* was also expressed in *Tmtc1-*negative, SLC14A1-negative medullary vessels (**Fig. S7A-B,** arrowheads). We traced the medullary *Esm1-*lineage in relation to SLC14A1 DVR using *Esm1*CreERT2; mT/mG as above. At E18.5 (induced at E17.5), GFP+ cells were found in SLC14A1-negative medullary vessels (**Fig. S7C**). When traced from P0 to P3, a portion of GFP+ cells had incorporated into the DVR, including some cells that connected SLC14A1-vessels and SLC14A1+ DVR capillaries (**Fig. S7D-D”**). By P15 (traced from P0), GFP+ cells in the inner medulla were primarily located in DVR capillaries (**Fig. S7E-E”**). Outer medullary GFP+ cells were variably in DVR, with numerous individually labeled GFP+ cells still present 15 days after labeling (**Fig. S7E-E’”**). Together, these results suggest that *Esm1*+ cells in the medulla contribute to DVR in a centripetal manner.

## Discussion

### A transcriptional atlas of the mouse embryonic kidney endothelium

Organotypic heterogeneity of vascular endothelial cells (ECs) has been of major interest in recent years.^47^ Local factors, including chemical or mechanical cues, mold the transcriptional identity of ECs into recognizable vessel subtypes that carry out tissue specific functions. Here, we present a developmental atlas of embryonic renal ECs, identifying distinct transcriptional signatures for proliferative and angiogenic vessels that contribute to proper renal vascularization.

Previous efforts to map the renal vasculature, including our own, have had limited success in producing validated specific marker genes for certain kidney vessel types. Genes such as *Plvap, Aplnr, Emcn* have been posited as useful *in silico* marker genes of AVR, progenitor, and glomerular ECs respectively,^9,48,49^ but *in vivo* have been shown to be highly non-specific. While differentially expressed genes from previous single cell studies provide useful transcriptional signatures with high sensitivity, we believe that our validated markers such as *Sgk1, Chrm3, Tmtc1, Fbln2, Jam2* and *Ccdc3* with restricted expression to certain vessel beds will be invaluable to researchers studying renal vasculature *in vivo* without single cell methodology.

An important caveat to this study is that we failed to capture ECs of large veins and peritubular capillaries within our single nuclear dataset and marker genes. We speculate this is due to low expression of the *Flk1::*H2B-eYFP reporter in these endothelial subpopulations (**Fig. S1**) or to a relative paucity of PTCs at E18.5 (especially distal tubules) compared to other developmental vessel types. Using *Esm1* lineage tracing, we found that vessels of the NZP surprisingly do not contribute to the PTCs. How the PTCs develop, perhaps via direct contribution of the SNZP, will be of great interest.

### Ccdc3 specifically marks proliferative vascular progenitors of the sub-nephrogenic plexus

This study provides several novel insights into *in vivo* renal vascular development. We previously described the dense vascular plexus underlying the NPCs as early as E13.5, which we term the ‘sub-nephrogenic plexus’. Barry et al. identified an *Aplnr*-enriched progenitor population by pseudotime analysis of single cell endothelial data.^9^ Here we validate and build upon these findings. Importantly, our new marker gene *Ccdc3* is specifically expressed in sub-nephrogenic plexus ECs and not in terminal vessel types, whereas *Aplnr* remains expressed in mature kidney vessels. Co-enrichment of classical vein markers such as *Nr2f2* and direct physical connectivity between the sub-nephrogenic plexus and lumenized veins suggest a venular identity, supporting previous pseudotime analysis showing vascular progenitors arising directly from veins.^9^

Importantly, we demonstrate the proliferative nature of the sub-nephrogenic plexus, using both pHH3 staining combined with *Ccdc3* expression as well as *Fucci2a* genetic reporter labeling. This analysis reveals a subtle but important distinction from other cell types in the kidney. While the stromal, nephron, and ureteric lineages all undergo proliferation in the NZ at the periphery of the cortex, we find that the *Ccdc3*+ sub-nephrogenic vessels, not the NZ vessels, are the most highly proliferative during development, suggesting potential EC-specific, zonated regulation of proliferation. Future studies into the origins, regulation, and ultimate resolution of the sub-nephrogenic plexus will be crucial to understanding formation of all kidney vasculature.

### Tip cells in the nephrogenic zone contribute to glomerular capillaries

In this study, we identify a distinct cluster of angiogenic, filopodia-bearing, *Esm1*+ tip cells in the E18.5 kidney, which have not been previously characterized *in silico* or *in vivo*. Using *Esm1*CreERT2 tracing, we visualize NZ tip cells and map their integration into cortical distal arteries and glomeruli. We find different categories of tip cells that localize to distinct parts of the NZ, potentially indicating differing trajectories. Peripheral tip cells extend multiple projections between NPC caps to potentially initiate the next round of iterative renewal of the NZP.^10^ Munro and colleagues previously described a continuous path between arteries and these bifurcating vessels, indicating that these tip cells may contribute to arterial vasculature as we see in lineage tracing experiments. Alternatively, tip cells between the ureteric bud (UB) and NPC cap surround early renal vesicles with filopodia. These intricate cells have not been previously described as part of the NZ vasculature but likely contribute to glomerular vascularization.

We previously showed that the developing nephron is enwrapped by a “net” of vessels starting at the renal vesicle stage.^8^ Here, we show that this net contains long-lasting tip cells that continue to extend filopodia towards epithelial components until the S-shaped body (SSB) stage. Tip cells are found in different locations surrounding SSBs, revealing new potential contributions to glomerular vasculature. Sprouting tip cells within the glomerular cleft form arteriolar vasculature, in coordination with a separate potential sprouting event from endothelial cells on the opposite side of the podocytes. We observe stromal cells expressing NG2, a marker of mural cells including vascular smooth muscle and pericytes, surrounding a portion of the developing arterioles within the glomerular cleft, providing further evidence that *Esm1-*lineage cells within the cleft contribute to arterioles.

Importantly, lineage tracing data suggests that all SSB-associated tip cells, including those outside the glomerular cleft, contribute to glomerular or arterial vasculature. This is surprising, as logically endothelial cells near segments of the SSB that give rise to cortical tubules would contribute to peritubular capillaries. Instead, we find that tip cells outside of the glomerular cleft reach ‘behind’ the SSB to incorporate into a vascular net, which later associates tightly with arteriolar vasculature. We posit that this vascular plexus contributes individual cells to form glomerular loops. In addition, we observe NG2+ stromal cells interspersed between endothelial cells of the vascular net. As NG2 is expressed in a variety of stromal cells including smooth muscle and mesangial cells, the exact identity of these vascular associated cells in the mature glomerulus remains unclear. In sum, rather than one sprouting vessel entering the glomerular cleft and looping to form the entire glomerular and arteriolar vasculature, we propose at least three different sources of endothelial cells that specifically contribute to inflow, outflow, and capillary tufts.

In accordance with this model, the *Esm1* lineage contributes to glomeruli in a distinctly mosaic, non-stereotyped fashion, even when traced to P15 to exclude the possibility of lingering tamoxifen induction. Labeling of one arteriole, as well as individually labeled GFP+ cells within the glomerular capillaries, support our model in which cells within the glomerular cleft contribute to half of the arteriolar vasculature, while cells in the net ‘behind’ the SSB contribute individual ECs in an additive fashion.

Importantly, we characterize the multi-clonal nature of glomerular vasculature as arising from distinct tip cells, rather than one sprouting event giving rise to an entire glomerular vasculature. At either stage, the labeling of isolated cells in multiple locations around the developing glomerulus suggests a distinct clonal nature (i.e. cells within the glomerular cleft and the vascular net are not always labeled in conjunction). To demonstrate this incontrovertibly, we used a Rainbow reporter to show that glomerular vascular development frequently involves multiple tip cell clones contributing to visually distinct parts of the endothelium.

These observations are based on fixed timepoint analysis, so their validity and biological implications will require future intravital imaging or *ex vivo* studies. However, the preponderance of the evidence supports multi-clonal mechanism of glomerular vascularization, involving tip cell contributions from ECs found in various locations around the developing nephron.

Taken together, these results provide new insight into the cellular mechanism of glomerular vascularization. We find angiogenic tip cells associated with the developing nephron from the earliest stages of epithelial development. These tip cells sprout into the glomerular cleft, but importantly, a single sprouting vessel does not account for the entirety of a glomerular vasculature as previously thought,^50^ as GFP+ cells join with non-GFP+ endothelium across the glomerular cleft to form arteriolar vasculature (**Fig. 6N**). Importantly, we find that tip cells associated with other portions of the SSB also contribute to glomerular vasculature in a clonal, mosaic fashion, with no significant contribution to other cortical capillaries. These cells incorporate into a dense vascular net that forms as the SSB progresses, which likely contributes individual cells to the glomerular capillaries, as opposed to previous models of progressive looping of an endothelial vessel within the glomerular cleft.

### Proximal and distal renal arterial development is differentially regulated

We and others have previously described the ramifying branching pattern of the renal arterial tree,^8,51^ but the details of arterial formation and connection to glomerular vasculature remain unclear. Here, we show multiple previously unknown molecular distinctions between the proximal (large) arteries and the distal (small) arteries and arterioles. At E18.5, distal and proximal arterial ECs cluster distinctly. In addition, our lineage tracing suggests that distal arteries originate from cortical tip cells. Whether this difference in cellular origin is related to transcriptional distinctions remains to be seen. Importantly, *Esm1* is not expressed in proximal arteries at E18.5, suggesting that distal arteries are derived from cortical tip cells. These results mirror previous studies of the mouse retina showing that *Esm1+* tip cells give rise to arteries.^52^ As with our own findings regarding glomerular development, the exact steps of distal artery formation are difficult to ascertain without time-lapse imaging. In addition, how tip cell-mediated distal arterial development and glomerular vascularization coordinate remains unclear.

Finally, our Fucci2a analysis suggests differential cell cycle regulation along the arterial tree. Chavkin and colleagues recently showed in the mouse retina that late G1 arrest, marked by FucciRed enrichment, in ECs drove TGFβ1-mediated arterial gene expression.^35^ Intriguingly, in both E18.5 and P15 kidneys, proximal arteries, but not distal arteries, were enriched for FucciRed+ nuclei, indicating that distal arterial specification in the kidney may rely on pathways independent of cell cycle arrest. Future studies will be needed to elucidate the mechanistic underpinnings of these differences between proximal and distal arteries, and our identification of novel markers of the two artery types will be invaluable.

### Novel insights into vasa recta biology

The development of the vasa recta, despite being one of the most specialized endothelial features of the kidney, remains largely unexplored. Here, we show that *Tmtc1* expression is specific to the AVR, without cortical or DVR expression, which will aid characterization of the medullary vasculature along with DVR markers such as *Slc14a1*. In addition, we uncover novel biological features of the vasa recta, suggesting interesting underlying mechanisms. Genes such as *Clec14a,* which are expressed in all vasa recta capillaries, suggest overarching transcriptional differences between medullary and cortical endothelium. In addition, at both E18.5 and P15 we observed widespread medullary G1 arrest marked by FucciRed enrichment in vasa recta. Whether these medullary specific signatures drive vasa recta development or are an adaptive response to the local acidic, hypertonic, or hypoxic environment remains to be answered.

We also provide new insights into the growth of the vasa recta, both during embryonic development and post-natally. Medullary *Esm1+* lineage traced ECs pulsed at P0 originate in vessels outside of either ascending or descending vasa recta, and yet they eventually contribute to the DVR in a centripetal manner. In addition, both *Esm1* expression and proliferative ECs are restricted to the outer medulla at P15, suggesting the presence of a pool of proliferative ECs that may support longitudinal growth of the entire vasa recta, including via *Esm1+* cell-driven DVR contribution. Further investigation as to whether these *Esm1+* cells represent true tip cells and their relation to proliferating ECs in the outer medulla will further clarify the development of the vasa recta post-natally.

### Utility of an embryonic kidney renal atlas

High resolution characterization of regionalized blood vessels in the kidney will provide tools for studying both kidney disease, organ replacement, and regeneration. We recently reported the lack of vasculature and failed perfusion of classic iPSC derived kidney organoids.^53^ Multiple strategies have attempted to address this challenging problem to varying degrees of success.^11,54^ Importantly, we believe our developmental endothelial atlas will improve our ability to evaluate organoid vasculature. While assessing maturity of nephron and ureteric cell types is possible due to in-depth molecular characterization of epithelial development,^41^ evaluation of ECs has thus far been limited to terminal (mature) transcriptional signatures. Our characterization of progenitor, transitional, and angiogenic ECs will help benchmark progress of organoid vascularization efforts and identify required cues for proper renal vascular growth.

A renal vascular transcriptional atlas will also aid characterization of developmental vascular phenotypes in functional genetic studies. In the present study, we utilized markers of the proliferative sub-nephrogenic and angiogenic NZP to characterize ectopic vasculature in netrin 1 knockout mice. Using our new tools, we observe an increase in ECs expressing neither *Esm1* nor *Ccdc3* at the kidney surface these mutants, indicating a loss of normal developmental EC states. This could be associated with alterations in EC proliferation or tip cell behavior, setting up future avenues of investigation. Applying the developmental markers and insights from this study will be of great interest in other models of altered kidney vascular development.

### Conclusions

In this study, we provide a unified view of the developing kidney vasculature using single-nucleus sequencing and *in vivo* characterization. We identify molecular zonation of the developing vasculature, and provide novel, specific marker genes for multiple renal vessel types. We show unique biological characteristics such as cell cycle state within vessel types such as the glomeruli, arteries, and vasa recta. Finally, we extensively characterize novel, developmentally restricted, vascular beds. We detail a proliferative, venous *sub-nephrogenic plexus* (SNZP) and a connected, angiogenic *nephrogenic zone plexus* (NZP). We perform detailed lineage tracing to show that tip cells in the nephrogenic zone contribute in a multi-clonal manner to glomeruli and arteries, providing extensive evidence to support a new model of nephron vascularization. We hope this study will be a valuable resource to researchers interested in kidney, developmental, and vascular biology, and will inform and direct future studies into renal vascular development.

## Methods

### Mouse Handling

Experiments were performed in accordance with protocols approved by the UT Southwestern Medical Center IACUC, Northwestern University School of Medicine or University of Virginia School of Medicine, depending on the location the strain was housed. Cd1, *Flk1::H2B-eYFP, Cspg4-mEGFP, Foxd1Cre; Ntn1^ff^* mice were all house at UT-Southwestern. *Cdh5CreERT2; Fucci* mice were housed at the University of Virginia School of Medicine. *Esm1CreERT2; Rainbow* and *Esm1CreERT2; mTmG* mice were housed at Northwestern University School of Medicine.

All mouse embryos and postnatal pups were used without sex identification (mixed sexes). For a complete list of all strains used in this study, please see the key strains table. For all mice used in the study, food and water were given ad libitum. No procedures had been done to the mice prior to use in this study. Prior to study, mice were housed with same sex littermates in groups of 3-5, in accordance with IACUC policy.

For embryonic studies (E15.5 - E18.5), timed pregnancies were induced by placing one male with one to two females of the appropriate genotype in a cage overnight. Plug checking was each morning to check for pregnancy. Upon dissection, embryos were staged by checking for various morphological landmarks (snout morphology, paw morphology) to verify appropriate developmental stage. For postnatal studies (P3-P15), postnatal pups were left in the same cage with their mothers (in accordance with IACUC policy). P0 is defined as the date of birth.

### Single Nuclear Isolation and Sequencing

#### Nuclei Collection

Single nuclear isolation and sequencing was performed in accordance with previously published protocols. In brief, an E18.5 Flk1::H2B-eYFP pregnant dam was dissected, and embryos placed on ice. We then sorted whole embryos under an epifluorescent scope for visible YFP expression. Both kidneys were then removed from the YFP-positive embryos by manual dissection. For the single nuclear data presented in this study, kidneys from 5 YFP positive embryos (a total of 10 kidneys, from littermate embryos) were pooled and then minced into small cubes (approximately 1mm^3^). Kidney pieces were placed in a precooled Eppendorf with 1mL of nuclear lysis buffer (Nuclei EZ Lysis Buffer, Sigma N-3408; 1 tablet of cOmplete ULTRA mini tablets, Roche 05 892 791 001). This suspension was then transferred to a Dounce grinder (Kimble Chase, KT885300-0002) with another milliliter of the nuclear lysis buffer was dounced with the loose pestle approximately 10 times. This suspension was then passed through a 200µm cell strainer (Puri select, 43-50200), and then further dounced with the tight pestle approximately 10 times with an additional 2mL of nuclear lysis buffer. The homogenate was then passed through a 40µm cell strainer (Puri select, 43-50040), and the run through was collected in a 15mL Falcon tube and centrifuged for 5 minutes at 500g at 4°C. The supernatant was discarded, and the pellet was resuspended in 4mL of nuclear lysis buffer 2 (Nuclei EZ Lysis Buffer; RNasin Plus, 1:1000, Promega N2615; SUPERaseIN, 1:1000, Thermo Fischer, AM2696). The sample was then incubated on ice for 5 minutes, and centrifuged again or 5 minutes at 500g at 4°C. The supernatant was then discarded, and then pellet was resuspended in nuclear suspension buffer (2% Bovine Serum Albumin in PBS, with SUPERaseIN, 1:1000). We then took 10µL of the nuclei suspension and mixed with Trypan Blue to evaluate nuclei integrity and number on a hemocytometer. We estimated that we obtained approximately 6 million total nuclei from the kidneys. This bulk suspension was then taken to the Children’s Research Institute (CRI) Flow cytometry core, and YFP positive cells were then sorted using FACSAria II in accordance with their standard protocols. After sorting, we recounted nuclei with a hemocytometer as above, as estimated that had obtained 100,000 YFP endothelial cells in total.

#### Sequencing

These nuclei were then taken to the Next Generation Sequencing Core and used for sequencing according to their standard protocols using the 10X Genomics Next GEM Single cell 3’ Reagent Kit v3.1. In brief, single nuclei suspensions are washed in 1X PBS (calcium and magnesium free) containing 0.04% weight/volume BSA (400 μg/ml) and brought to a concentration of ∼700-1200 cells/μl. This is accomplished by staining 10 μl of the single cell suspension with Trypan blue and reading on the Countess™ II Automated Cell Counter (ThermoFisher). The appropriate volume of cells is loaded with Single Cell 3’ Gel Beads into a Next GEM Chip G and run on the Chromium™ Controller. GEM emulsions are incubated and then broken. Silane magnetic beads are used to clean up GEM reaction mixture. Read one primer sequence is added during incubation and full-length, barcoded cDNA is then amplified by PCR after cleanup. Sample size is checked on the Agilent Tapestation 4200 using the DNAHS 5000 tape and concentration determined by the Qubit 4.0 Fluorimeter from ThermoFisher using the DNA HS assay. Samples are enzymatically fragmented and undergo size selection before proceeding to library construction. During library preparation, Read 2 primer sequence, sample index and both Illumina adapter sequences are added. Subsequently, samples are cleaned up using Ampure XP beads and post library preparation quality control is performed using the DNA 1000 tape on the Agilent Tapestation 4200. Final concentration is ascertained using the Qubit DNA HS assay. Samples are loaded at 1.6 pM and run on the Illumina NextSeq500 High Output Flowcell using V2.5 chemistry. Run configuration is 28x98x8.

#### Single-nucleus RNA-seq preprocessing

Filtered feature-barcode matrices were generating by running the CellRanger count pipeline. Nuclei were called from empty droplets by testing for deviation of the expression profile for each cell from the ambient RNA pool.^55^ Nuclei with low numbers of detected genes or library sizes, as defined as a deviation greater than three median absolute deviations (MADs) below the median, were excluded from further analysis. Nuclei with large mitochondrial proportions, i.e., more than 3 mean-absolute deviations above the median, with a minimum difference between the threshold 0f 0.5% between the threshold for exclusion and the median mitochondrial transcript count, were removed. Nuclei were pre-clustered and a deconvolution method was applied to compute size factors for all cells^56^ and normalized log-expression values were calculated.

#### Single-nucleus RNA-seq analysis

Cells were clustered by building a shared nearest neighbor graph^57^ on the normalized expression data and executing the Louvain algorithm. Diffusion maps were calculated using an adaption of the algorithm presented in Haghverdi et al.^58^ Normalized expression data was imputed applying Markov Affinity-based Graph Imputation of Cells (MAGIC)^59^ to the previously constructed diffusion maps. Marker genes for each subset of cells under investigation were calculated by applying the EIGEN algorightm.^60^ Differential expression was measured by calculating the earth mover’s distance (Wasserstein-1) distance between the distribution of imputed expression of each gene supported by the subsets of cells being compared. Gene set enrichment analysis was performed for sets curated in MsigDb^61^ using with fgsea^62^ Bioconductor package with the earth-mover distance as the ranking statistic.

### RNAScope

All RNAScope experiments were carried out on fresh frozen tissue. To prepare tissue, samples were dissected in PBS, and then gently dabbed on a Kimwipe to remove excess liquid. Samples were then embedded in OCT (Tissue Tex, 4583) and flash frozen. Embedded blocks were sectioned on a Leica Cryostat at a sectioning thickness of 12 µm onto SuperFrost glass slides (Fisher Scientific, ***) and slides were stored at −80°C until assayed.

RNAscope Hiplex experiments were carried out using manufactures protocol using the RNAScope Hiplex v2 kit. Sections were placed in 4% PFA at room temperature for 1 hour, and then progressively dehydrated to 100% Ethanol. Samples were then permeabilized using Pro3 from the Hiplex v2 kit. For E15.5, E18.5 and P4 kidneys, samples were permeabilized for 15 minutes at room temperature. For P15 kidneys, samples were permeabilized for 30 minutes at room temperature. Probe hybridization was carried out at 40°C for 2 hours. After probe hybridization, samples were washed twice for 2 minutes in Hiplex wash buffer. We then added Amp1 for 30 minutes at 40°C, washed twice in Hiplex was buffer, and then repeated with Amp2 and Amp3. Fluorophore conjugation was performed for 15 minutes at 40°C, DAPI was added for 30 seconds and slides were subsequently mounted in Prolong Gold mounting media (P36934). RNA-Scope imaging was performed on either the Nikon CSU-W1 or Nikon CSU-W1 SoRa confocal microscopes at the Quantitative Light Microscopy Core at UTSW. After imaging, slides were soaked in 4x SSC until the coverslips fell off and then were placed in freshly prepared cleaving solution. Slides were cleaved for 15 minutes and then washed with 0.5% PBST at room temperature. This process was then repeated (a total of 2 cleaving steps for each round of fluorophores, 30 minutes of cleaving total). The next fluorophore conjugation step was carried, at 15 minutes at 40°C, and DAPI and slide mounting was performed as above. The same section was imaged on the confocal as above. This process was repeated until all four fluorophore sets were images.

After all probe sets were imaged, we then cleaved the last round of fluorophores off in preparation for immunofluorescence. Cleaving was carried out as above. Post cleaving, we permeabilized slides 0.3% Triton in PBS for 10 minutes. Slides were blocked in Cas Block (Invitrogen) for approximately 1 hour. Primary antibodies were added overnight at 4 degrees Celsius, and then washed in PBS three times, 5 minutes per wash with gentle agitation. Secondary antibodies were added for 1-2 hours at room temperature. Slides were then mounted and images as above.

To process RNAscope images, maximum intensity projections were created in ImageJ Fiji. These images were then loaded into the ACD Hiplex Registration software and registered such that we are able to overlay fluorophores from subsequent imaging rounds. Alternatively, some images were manually registered in ImageJ Fiji.

RNAscope signal corresponding to background and non-endothelial signal was removed as noted in figure legends from Fig. 1D and Fig. 2X (*Tmtc1* only). One of two methods was used to create a mask of *Pecam1* signal. For Fig. 1D, the Spots function in IMARIS was used to identify *Pecam1* RNAscope dots with the following parameters: Quality ****. Following Spot creation, Spot size on IMARIS viewer was adjusted to 5um (calculated to the nearest whole pixel) the mask was exported from the viewer. A ROI for the *Pecam1* mask was created in Fiji using Analyze Particles, selecting ‘vessels’ with at least two contiguous spots. Signal outside of the ROI was cleared.

For Fig. 2F, a *Pecam1* mask was created as follows. A 3um Gaussian blur was applied in Fiji and signal was thresholded with the following settings: A ROI was created using Analyze Particles in Fiji and signal outside of the ROI was cleared.

### Immunofluorescence (IF) staining on tissue sections

Unless otherwise noted, antibody staining was performed on fresh frozen cryosections. For parraffin sections: Tissue was fixed in 4% PFA overnight at 4 degrees Celsius. After dissection and fixation, tissue was dehydrated to 100% ethanol through a standard ethanol gradient. Tissue was cleared with xylene twice for 10 minutes each (15 minutes x2 for P15 tissue), and incubated in 50/50 xylene/Paraplast X-TRA (Leica) for 10 minutes in a 55-60° bead bath. Tissue was incubated in Paraplast X-TRA for up to 18 hours, replacing with fresh Paraplast at least three times and incubating overnight in a 55-60° oven. Tissue was then positioned and embedded into an embedding mold, and sectioned at 10um on a Leica microtome onto SUPERFROST glass slides (Fisherbrand).

After drying sectioned slides overnight, slides were baked in a 60° oven for 10 minutes. Slides were deparaffinized with xylene twice for 5 minutes, and rehydrated through an ethanol gradient. Slides were dipped in de-ionized water, washed in PBS three times for 5 minutes, and permeabilized with 0.3% Triton-X in PBS for 10 minutes with gentle rocking. Slides were then subjected to heat-induced antigen retrieval in R-Buffer A (Electron Microscopy Sciences) for nuclear antigens or R-Buffer B (Electron Microscopy Sciences) for cytosolic or membrane-bound antigens using a Retriever 2100 (Electron Microscopy Sciences). After cooling to room temperature, slides were dipped in PBS and blocked in CAS-Block (Invitrogen) for at least one hour. Slides were incubated in primary antibody mixture overnight at 4°, diluted in CAS-Block, in a humidity chamber. Slides were washed with PBS for 5 minutes three times the following day and incubated in secondary antibody mixture for 1.5 hours at room temperature. Slides were then washed three times in PBS for 5 minutes, and incubated in a blood lysis solution (CuSO4, NH4Cl, pH 5) for 15 minutes. Slides were incubated in de-ionized water for 5 minutes to lyse blood cells and washed in PBS two times for 5 minutes each. Slides were then mounted in DAPI-Fluoromount G (Southern Biotech) and imaged on Nikon CSU-W1 or Nikon CSU-W1 SoRa confocal microscopes at the Quantitative Light Microscopy Core at UTSW.

### Whole mount immunofluorescence (WMIF) on vibratome-cut sections

After dissection and fixation, tissue was embedded in 2.5-5% low melting point agarose (Sigma). 100µm sections were cut using a Leica vibratome, with speed .3mm/sec and amplitude 1mm. Sections were stored in glass vials or in 48-well plates to maintain positional information for selecting sections near the periphery of the kidney (edge sections). Staining was done in 96-well plates or 48-well plates for P15 tissue with gentle nutation. Sections were permeabilized with 1% Triton-X 100 in PBS for tissue younger than P4 and 3% Triton-X 100 in PBS for P15 tissue for 1.5-2 hours at room temperature. Sections were blocked in CAS-Block for at least 1 hour at room temperature, and incubated in primary antibody mixture overnight at 4°. Sections were washed in PBS for at least 5 hours, replacing with fresh PBS every hour, and were incubated in secondary antibody mixture overnight at 4°. Sections were washed in PBS at room temperature for at least 3 hours, replacing with fresh PBS once every 30 minutes. Sections were placed on SUPERFROST glass slides and cleared in a bubble of Rapiclear 1.47 (Sujin) for at least 30 minutes. Slides were mounted in Rapiclear and imaged on Nikon CSU-W1 or Nikon CSU-W1 SoRa confocal microscopes at the Quantitative Light Microscopy Core at UTSW.

#### *Esm1CreERT2* lineage tracing

*Esm1* lineage tracing was performed by crossing *Esm1*CreERT2 containing mice (male or female) to mice containing the *Rosa26::mT/mG* or *Rosa26::Rainbow* reporter alleles. For short-pulse lineage tracing designed to label *Esm1*-expressing cells, timed pregnancies were set up and pregnant dams were gavaged with 4-hydroxytamoxifen (4-OHT) at E17.5. Dams were sacrificed and embryonic kidneys were retrieved at E18.5. For long term lineage tracing designed to trace *Esm1+* cells throughout development, pregnant dams were monitored for date of birth, and pups were pipette-fed 4-OHT at P0. Pups were sacrificed at either P3 or P15 (mT/mG lineage tracing) or P4 (Rainbow lineage tracing). For embryonic timepoints, pregnant females were orally gavaged with 1mg 4-OHT dissolved in acetone and sunflower seed oil. Upon sacrifice, kidneys from all embryos in the litter were harvested for dehydration and genotyping. For postnatal timepoints, pups were orally gavaged with 5 uL of either 6 mg/mL or 20 mg/mL 4-OHT dissolved in ethanol and sunflower seed oil; whole litters of mixed males and females were used for experiments

#### *Fucci2a* endothelial cell cycle labeling

Endothelial cell cycle reporter labeling was performed by crossing *Cdh5*CreERT2 containing mice to mice containing the Rosa26::Fucci2a allele. For E18.5 kidneys, timed pregnancies were set up and reporter expression was induced at E15.5 with tamoxifen. For P15 kidneys, pups were injected with tamoxifen at P4. Tamoxifen (Sigma Cat# T5648) was resuspended in 10% EtOH and 90% Corn oil (Sigma Cat# C8267) at 4 mg/mL, and 25 μL was injected per pup at P4. The pregnant mouse was injected with 10 mg/mL (250 μL/mouse) for the embryonic timepoint (E18.5).

### Data processing and Statistics

#### RNAscope quantification

For Figure 3D, quantification of *Esm1* or *Ccdc3* expressing cells within the nephrogenic zone was performed as follows. Endothelial nuclei were visually identified within the nephrogenic zone using *Pecam1* RNAscope signal. *Esm1* or *Ccdc3* positivity was determined by at least 3 specific RNAscope dots.

#### Fucci2a quantification

All quantifications were performed on maximum intensity projections of 100um sections through the center of the kidney (medulla is visible). Each data point represents a cortical ROIs selected using LTL immunofluorescence. Vessels of the nephrogenic zoneplexus, sub-nephrogenic zone plexus, and peritubular/glomerular capillaries were identified visually using EMCN and LTL immunofluorescence. Total number of FucciRed and FucciGreen+ nuclei in each zone were counted.

### Key Resources Table

| REAGENT or RESOURCE | SOURCE | IDENTIFIER |
| --- | --- | --- |
| Antibodies |  |  |
| Goat polyclonal anti-VECAD (1:100) | R&D Systems | AF1002 |
| Rat polyclonal anti-EMCN (1:100) | Santa Cruz | sc-65495 |
| *Lotus tetragonolobus* lectin, biotin conjugated (1:100) | VectorLabs | B-1325 |
| Rabbit monoclonal anti-ERG (1:100, paraffin) | abcam | ab92513 |
| Goat polyclonal anti-YFP (1:100, paraffin) | MyBioSource | MBS448123 |
| Mouse monoclonal anti-COUPTFII (1:100, paraffin) | R&D Systems | PP-H7147-00 |
| Goat polyclonal anti-PODXL (1:100) | R&D Systems | AF1556 |
| Rabbit polyclonal anti-SLC14A1 (1:100) | Invitrogen | PA5-110377 |
| Rabbit polyclonal anti-CX40 (1:100) | Invitrogen | 36-5000 |
| Rabbit polyclonal anti-PAX2 (1:100) | Proteintech |  |
| Rabbit monoclonal anti-GFP (1:100) | Invitrogen | G10362 |
| Goat polyclonal anti-tdTomato (1:100) | MyBioSource | MBS448092 |
| Rat monoclonal anti-PECAM1 (1:100) | BD Biosciences | 550274 |
| Rabbit polyclonal anti-pHH3 (1:100) | Millipore Sigma | 06-570 |
| Mouse monoclonal anti-Aqp1 (1:100) | Novus Biologicals | NB600-749 |
| Mouse monoclonal anti-aSMA-FITC (1:100) | Millipore Sigma | F3777-.2ML |
| Rabbit polyclonal anti-SIX2 (1:100) | Proteintech | 11562-1-AP |
| *Dolicho Biflorus* agglutinin (1:100) | VectorLabs | B-1035 |
| Mouse monoclonal anti-ECAD (1:100) | BD Biosciences | 610182 |
| Rabbit monoclonal anti-JAG1 (1:100) | Cell Signaling Technologies | 2620 |
| Goat polyclonal anti-SOX17 (1:100) | R&D Systems | AF1924 |
| Critical commercial assays |  |  |
| Manual Assay Hiplex RNAscope v2 | ACD | CAT NO. 324419 |
| Mm-*Pecam1*-NoxHs-T11 (1:50) | ACD | Cat No. 451501-T11 |
| Mm-*Esm1*-T3 (1:50) | ACD | Cat No. 411761-T3 |
| Mm-*Ccdc3*-T2 (1:50) | ACD | Cat No. 813291-T2 |
| Mm-*Jam2*-T9 (1:50) | ACD | Cat No. 467321-T9 |
| Mm-*Fbln2*-T12 (1:50) | ACD | Cat No. 447931-T12 |
| Mm-*Sgk1*-T8 (1:50) | ACD | Cat No. 434791-T8 |
| Mm-*Tmtc1*-T6 (1:50) | ACD | Cat No. 518221-T6 |
| Mm-*Chrm3*-T5 (1:50) | ACD | Cat No. 437701-T5 |
| Mm-*Clec14a*-NoxHs-T5 (1:50) | ACD | Cat No. 821161-T5 |
| Mm-*Sox17*-T5 (1:50) | ACD | Cat No. 493151-T5 |
| Mm-*Gja4*-T10 (1:50) | ACD | Cat No. 588591-T10 |
| Mm-*Unc5c*-T4 (1:50) | ACD | Cat No. 403211-T4 |
| Mm-*Neo1*-T5 (1:50) | ACD | Cat No. 884471-T5 |
| Mm-*Unc5b*-T6 (1:50) | ACD | Cat No. 884481-T6 |
| Mm-*Aplnr*-T6 (1:50) | ACD | Cat No. 436171-T6 |
| Deposited data |  |  |
| Raw and analyzed data | This paper | GEO: |
| Experimental models: Organisms/strains |  |  |
| Mouse: CD1: Crl:CD1(ICR) | Charles River Laboratories |  |
| Mouse: *Flk1*::H2B-eYFP: Tg(Kdr-H2B/EYFP)32Maba/J |  | RRID:IMSR_JAX:039608 |
| Mouse: *Cdh5*CreERT2: C57BL/6-Tg(Cdh5-cre/ERT2)1Rha | Taconic Biosciences | MGI:3848982 |
| Mouse: Fucci2a fl/fl: B6J;Cg-Gt(ROSA)26Sor<sup>tm1.1(CAG-Venus/GMNN,-Cherry/CDT1)Jkn/H</sup> | EMMA | EMMA:08395<br>MGI ID:6193738 |
| Mouse: *Esm1*CreERT2: Tg(Esm1-icre/ERT2)27Rha | Max Planck Institute for Molecular Medicine | MGI:7490347 |
| Mouse: mT/mG fl/fl: B6.129(Cg)-Gt(ROSA)26Sor<sup>tm4(ACTB-tdTomato,-EGFP)Luo/J</sup> | Jackson Laboratories | RRID:IMSR_JAX:007676 |
| Mouse: Rainbow fl/fl: Gt(ROSA)26Sor<sup>tm1(CAG-EGFP,-mCerulean,-mOrange,-mCherry)lw</sup> | Red-Horse, K. *et al.* 2010 | MGI:5441200 |
| Mouse: *Foxd1*-Cre: B6;129S4-*Foxd1*<sup>tm1(GFP/cre)Amc/J</sup> | Jackson Laboratories | RRID:IMSR_JAX:012463 |
| Mouse: *Ntn1* fl/fl: B6.129(SJL)-*Ntn1*<sup>tm1.1Tek/J</sup> | Jackson Laboratories | RRID:IMSR_JAX:028038 |
| Mouse: *Cspg4*-mEGFP: FVB.Cg-Tg(Cspg4-EGFP*)HDbe/J | Jackson Laboratories | RRID:IMSR_JAX:022735 |
| Software and algorithms |  |  |
| Fiji version 2.14.0/1.54f | Fiji | https://imagej.net/software/fiji/downloads |
| RNAScope Hplex Image Registration Software v2.1 | ACD |  |
| IMARIS 10.0 | Oxford Instruments |  |

## Supplemental Figure Captions

**Supplemental Figure 1.**
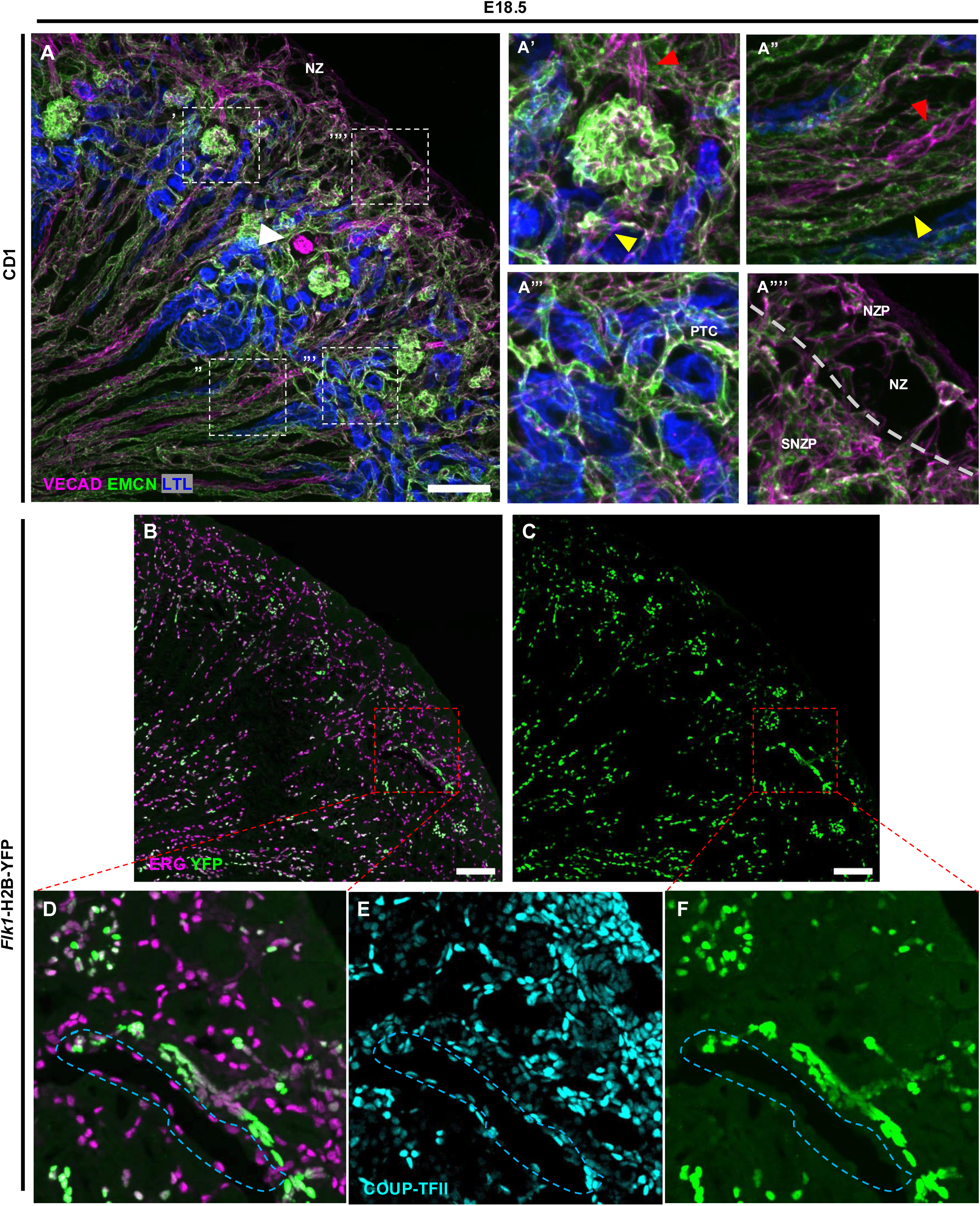
Differences in expression of broad vascular markers allows for visual identification of vessel types. (A) WMIF of 100μm E18.5 kidney section for VECAD, EMCN, and LTL. Arrowhead: VECAD+/EMCN- artery. (A’) Zoom of glomerular vessels and arterioles. Red, yellow arrowhead: Afferent (EMCN-) and efferent (EMCN+) arteriole, respectively. (A”) Zoom of vasa recta capillaries. Red, yellow arrowhead: Descending vasa recta (EMCN-) and ascending vasa recta (EMCN+), respectively. (A’”) Zoom of peritubular capillaries. (A’’’’) Zoom of periphery, showing nephrogenic zone plexus (NZP) and sub-nephrogenic plexus (SNZP). Dotted line: border of NZP. (B) ERG and YFP IF of E18.5 *Flk1*::H2B-eYFP kidney section. (C) YFP single channel view. (D) Inset of (B), showing corticomedullary artery and vein (dotted outline). (E) COUP-TFII IF of Inset from (B). (F) Inset of (C). Scale bar: 100µm (A-C).

**Supplemental Figure 2.**
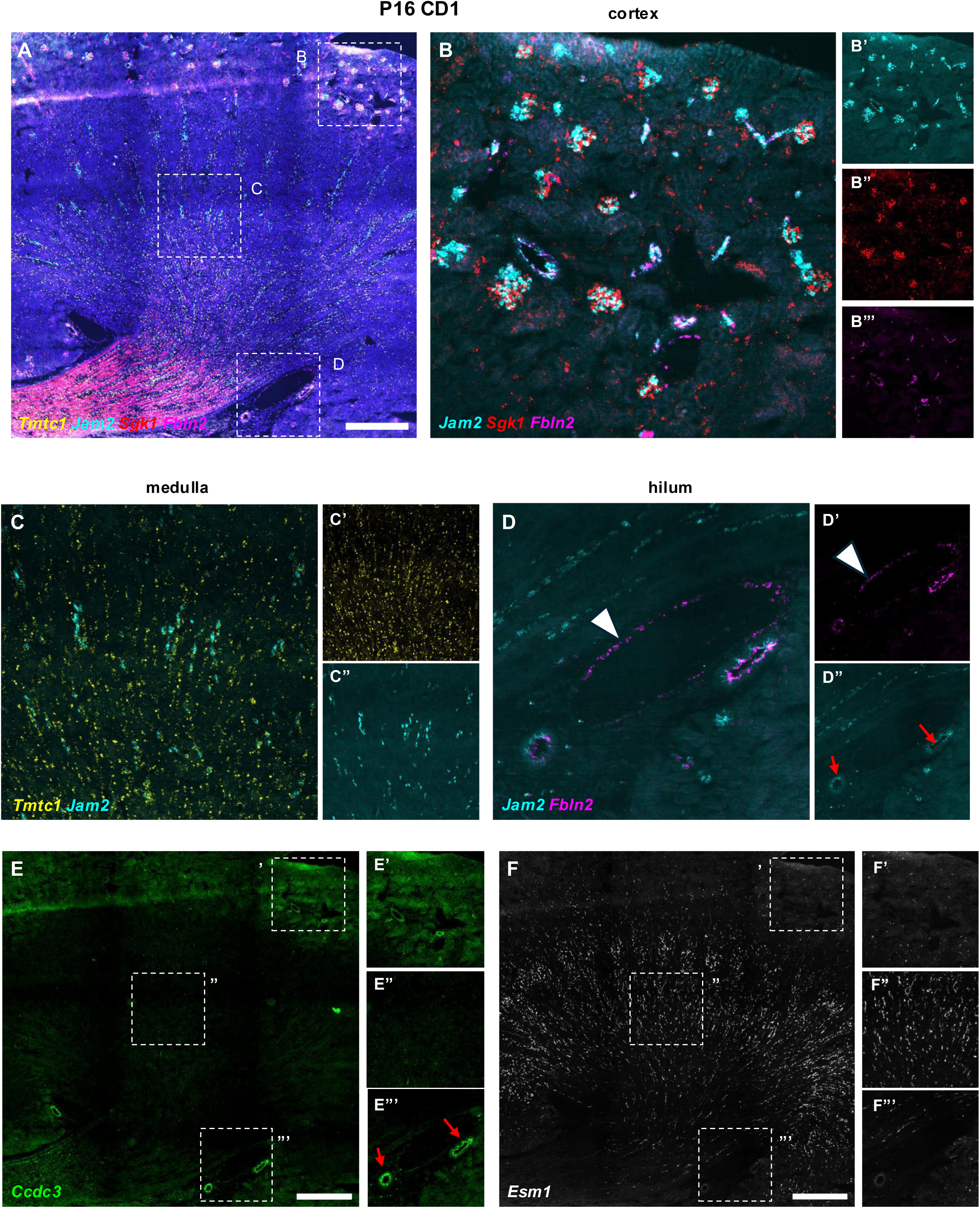
Markers of endothelial heterogeneity at E18.5 remain restricted after nephrogenesis. (A) *Tmtc1, Jam2, Fbln2,* and *Sgk1* RNAscope of P16 kidney section. (B) Zoom view of *Jam2, Fbln2,* and *Sgk1* in cortex. (B’-B”’) Single channel views of (B). (C) Zoom view of *Jam2* and *Tmtc1* in medulla. (C’-C”) Single channel views of (C). (D) Zoom view of *Jam2* and *Fbln2* in hilum. (D’-D”) Single channel views of (D). Arrowhead: vein with *Fbln2* expression. Red arrow: autofluorescence in arteries in *Jam2* channel. (E) *Ccdc3* RNAscope of section from (A). (E’-E”’) Zoom views of cortex, medulla, and hilum of (E). Arrow: autofluorescence in arteries. (F) *Esm1* RNAscope of section from (A). (F’-F”’) Zoom views of cortex, medulla, and hilum of (F). Scale bars: 500µm (A, E-F).

**Supplemental Figure 3.**
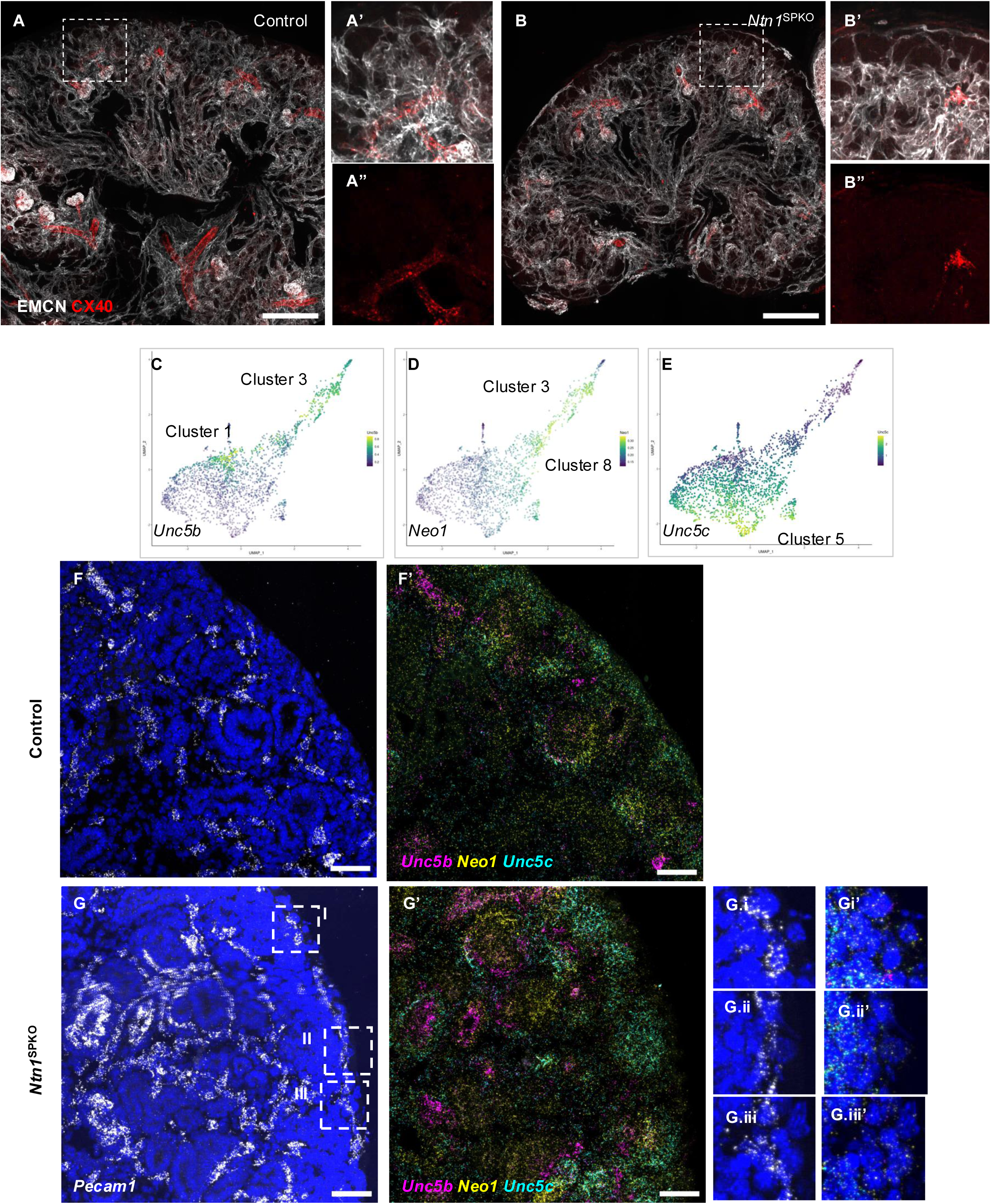
Ectopic vasculature in E15.5 netrin 1 knockouts does not display expression of either arterial genes or netrin 1 receptors. (A-B) WMIF of 100um E15.5 section of control and *Ntn1*^SPKO^ kidney for EMCN and CX40. (A’-B’) Zoom view of periphery of kidney. (A”-B”) CX40 single channel view of zoom. (C-E) UMAP of *Unc5b, Neo1,* and *Unc5c* expression in E18.5 single nuclear dataset. (F-G) *Pecam1* RNAscope of zoom view from **Fig. 3C** and **D**, respectively. (G’) *Unc5b, Neo1,* and *Unc5c* RNAscope of zoom view from (G). (G.I-G.III) Zoom view of *Pecam1* RNAscope of individual peripheral vessels, as in **Fig. 3C-I – Fig. 3C-III**. (G.I’-G.III’) Zoom views of (G’). Scale bars: 200µm (A-B), 50µm (F-G).

**Supplemental Figure 4.**
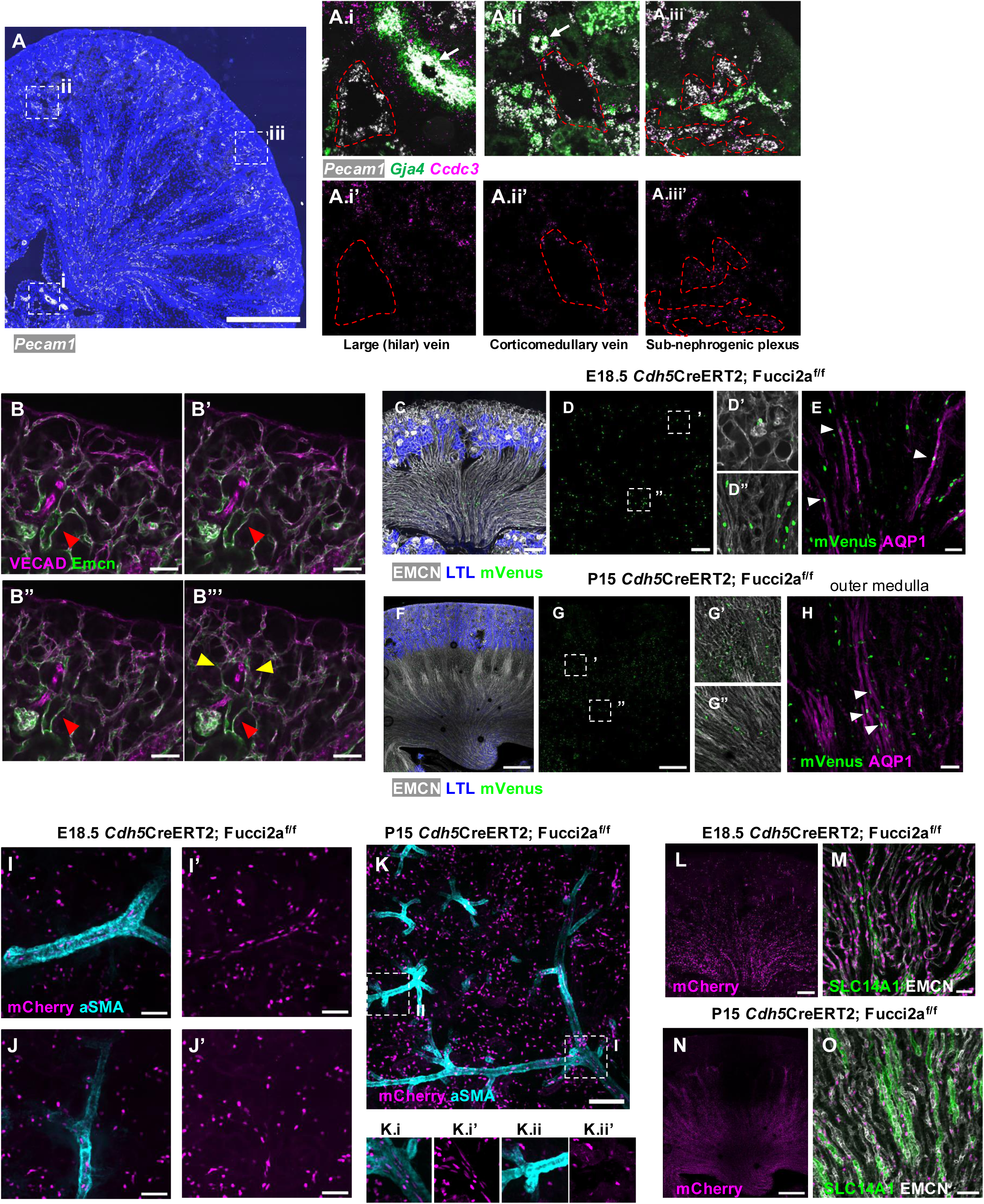
*Ccdc3* expression, venous lumens, and cell cycle regulation distinguish the sub-nephrogenic plexus from other renal vessel beds. (A) *Pecam1* RNAscope of E18.5 kidney section. (A.I-A.III) Zoom view of large and medium veins and sub-nephrogenic plexus, with RNAscope for *Ccdc3* and *Gja4*. Dotted line: venous lumen or sub-nephrogenic plexus vessels. (B-B”’) Serial z planes of WMIF of E18.5 kidney cortex for VECAD and EMCN. Red arrowhead: lumen of vein. Yellow arrowheads: Vessels connecting the sub-nephrogenic plexus with venous lumens. (C) WMIF of 100um section of E18.5 *Cdh5*CreERT2; Fucci2a^f/f^ kidney for EMCN, mVenus, and LTL. (D) mVenus only view of (C). (D’) Zoom view of EMCN and mVenus WMIF from (C), showing glomerulus (arrowhead). (D”) Zoom view of EMCN and mVenus WMIF from (C), showing medulla. (E) WMIF of 100um medullary section of E18.5 *Cdh5*CreERT2l Fucci2a^f/f^ kidney for mVenus and AQP1. Arrowheads: mVenus +/AQP1+ proliferative descending vasa recta cells. (F) WMIF of 100um section of P15 *Cdh5*CreERT2; Fucci2a^f/f^ kidney for EMCN, mVenus, and LTL. (G) mVenus only view of (F). (G’) Zoom view of EMCN and mVenus WMIF from (F), showing outer medulla. (G”) Zoom view as in (G’), showing inner medulla. (H) WMIF of 100um outer medullary section of P15 *Cdh5*CreERT2; Fucci2a^f/f^ kidney for mVenus and AQP1. Arrowheads: mVenus +/AQP1+ proliferating DVR cells. (I) WMIF of 100um section of E18.5 *Cdh5*CreERT2; Fucci2a^f/f^ kidney with proximal artery for mCherry and aSMA. (I’) mCherry only view of (I). (J) WMIF of 100um section of E18.5 *Cdh5*CreERT2; Fucci2a^f/f^ kidney with distal artery for mCherry and aSMA. (J’) mCherry only view of (J). (K) WMIF of 100um section of P15 *Cdh5*CreERT2; Fucci2a^f/f^ kidney for mCherry and aSMA. (K.I) Zoom view of proximal artery. (K.I’) mCherry only view of (K.I). (K.II) Zoom view of distal artery. (K.II’) mCherry only view of (K.II). (L) WMIF of 100um medullary section of E18.5 *Cdh5*CreERT2; Fucci2a^f/f^ kidney for mCherry. (M) WMIF of medulla of E18.5 *Cdh5*CreERT2; Fucci2a^f/f^ kidney for SLC14A1 and EMCN. (N) WMIF of 100um medullary section of P15 *Cdh5*CreERT2; Fucci2a^f/f^ kidney for mCherry. (O) IF of P15 *Cdh5*CreERT2; Fucci2a^f/f^ kidney showing medulla for SLC14A1 and EMCN. Scale bars: 500µm (A, F-G, N), 50µm (B, E, I-J, M, O), 200µm (C-D, L), 100µm (H, K).

**Supplemental Figure 5.**
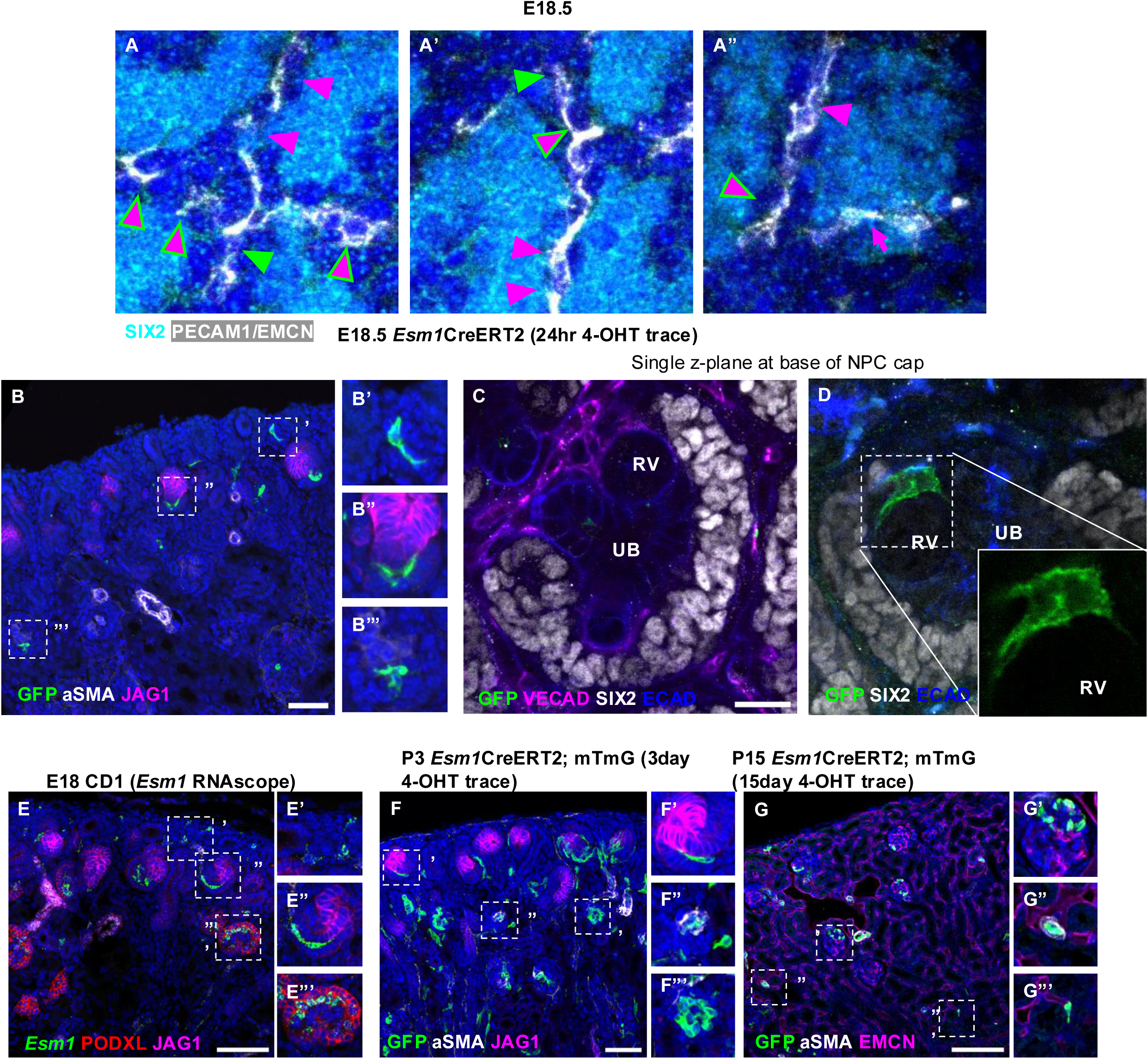
Details of tip cell location and fate in mouse kidney cortex. (A-A”) SIX2 and PECAM1/EMCN IF of zooms from **Fig. 5D’-D”’**. (B) IF of E18.5 *Esm1*CreERT2; mT/mG^f/f^ 24 hour lineage trace kidney for GFP, JAG1, and aSMA. (B’) Zoom view, showing peripheral GFP+ cell. (B”) Zoom view showing JAG1-associated GFP+ cell. (B”’) Zoom view, showing glomerular GFP+ cell. (C-D) Single z plane near base of NPC cap from **Fig. 5E-F**, respectively. RV, UB: Renal vesicle and ureteric bud. Inset, D: GFP+ cell with filopodia surrounding RV. (E) *Esm1* RNAscope with aSMA, JAG1 and PODXL IF on E18.5 kidney cortex. (E’-E”’) Insets of various vessel types. (F) GFP, aSMA, JAG1 IF on P3 *Esm1* lineage traced tissue. (F’-F”’) Inset of various vessel types. (G) GFP, aSMA, and EMCN IF on P15 *Esm1* lineage trace tissue. (G’-G”’) Insets of various vessel types. Scale bars: 50µm (B, F), 20µm (C-D), 100µm (E, G).

**Supplemental Figure 6.**
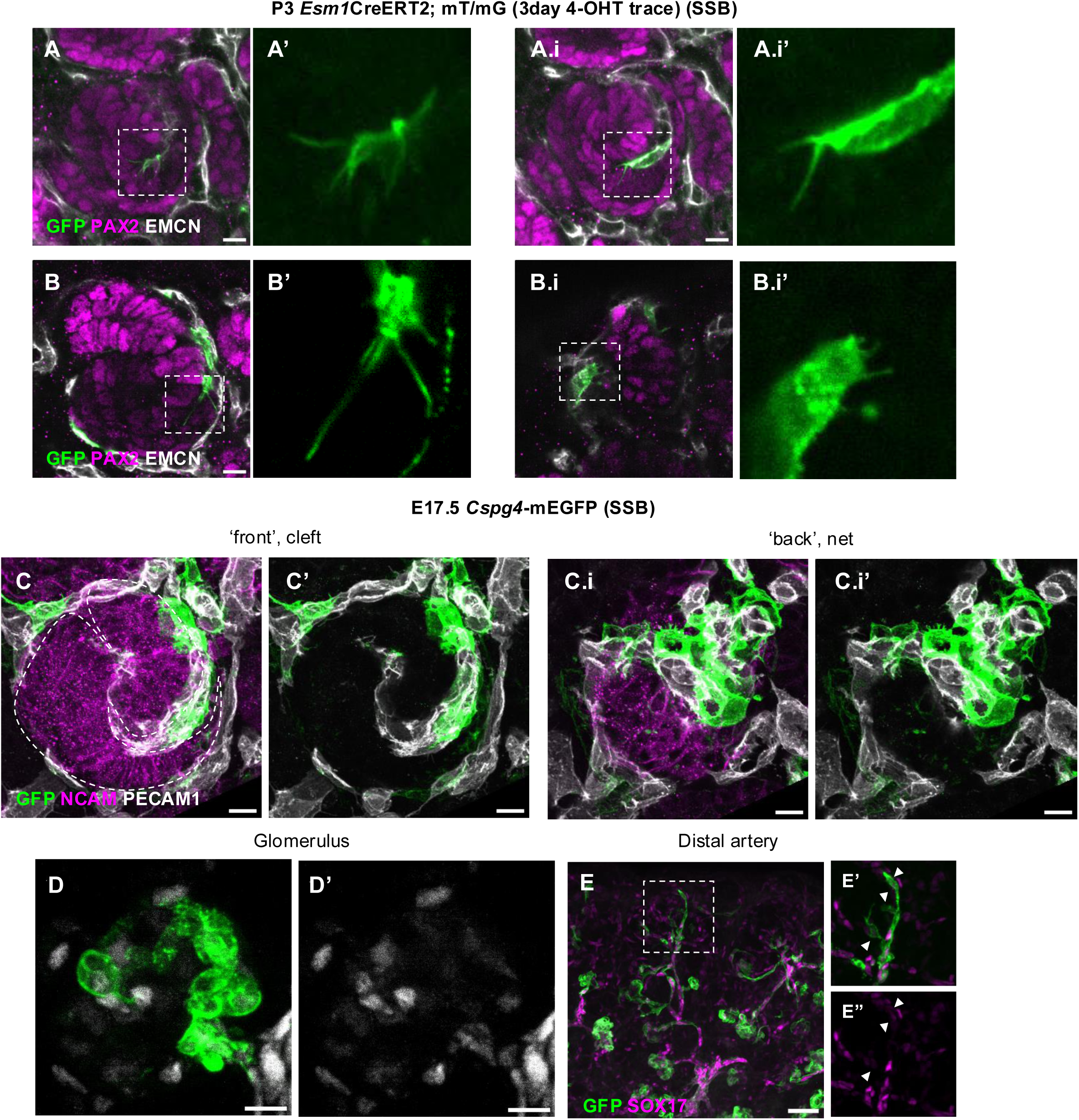
Additional details of tip cell morphology and contribution to glomerular and arterial vasculature. (A-A.i) Single z plane of **Fig. 6A**, showing tip cell filopodia. (A’-A.i’) Zoom view, GFP only of filopodia. (B) Single z plane of **Fig. 6B**, showing GFP+ cell in glomerular cleft. (B’) Zoom, GFP only of filopodia. (B.i) Single z plane of **Fig. 6B.i**, showing GFP+ cell behind SSB. (B.i’) Zoom, GFP only of filopodia. (C) WMIF of *Cspg4*-mEGFP, cortical section for GFP, NCAM, and PECAM1, showing front half of S-shaped body. (C’) GFP and PECAM1 only. (C.i) WMIF as in (C), showing back half of S-shaped body. (C.i’) GFP and PECAM1 only. (D) GFP and SOX17 WMIF of glomerulus from **Fig. 6G**. (D’) SOX17 only. (E) WMIF of 100um section of P3 *Esm1*CreERT2; mT/mG^f/f^ 3 day lineage trace kidney for GFP and SOX17. (E’) Zoom view of (E), showing distal artery formation. (E”) SOX17 only. Arrowheads: SOX17-low GFP+ cells. Scale bars: 10µm (A-D), 50µm (E).

**Supplemental Figure 7.**
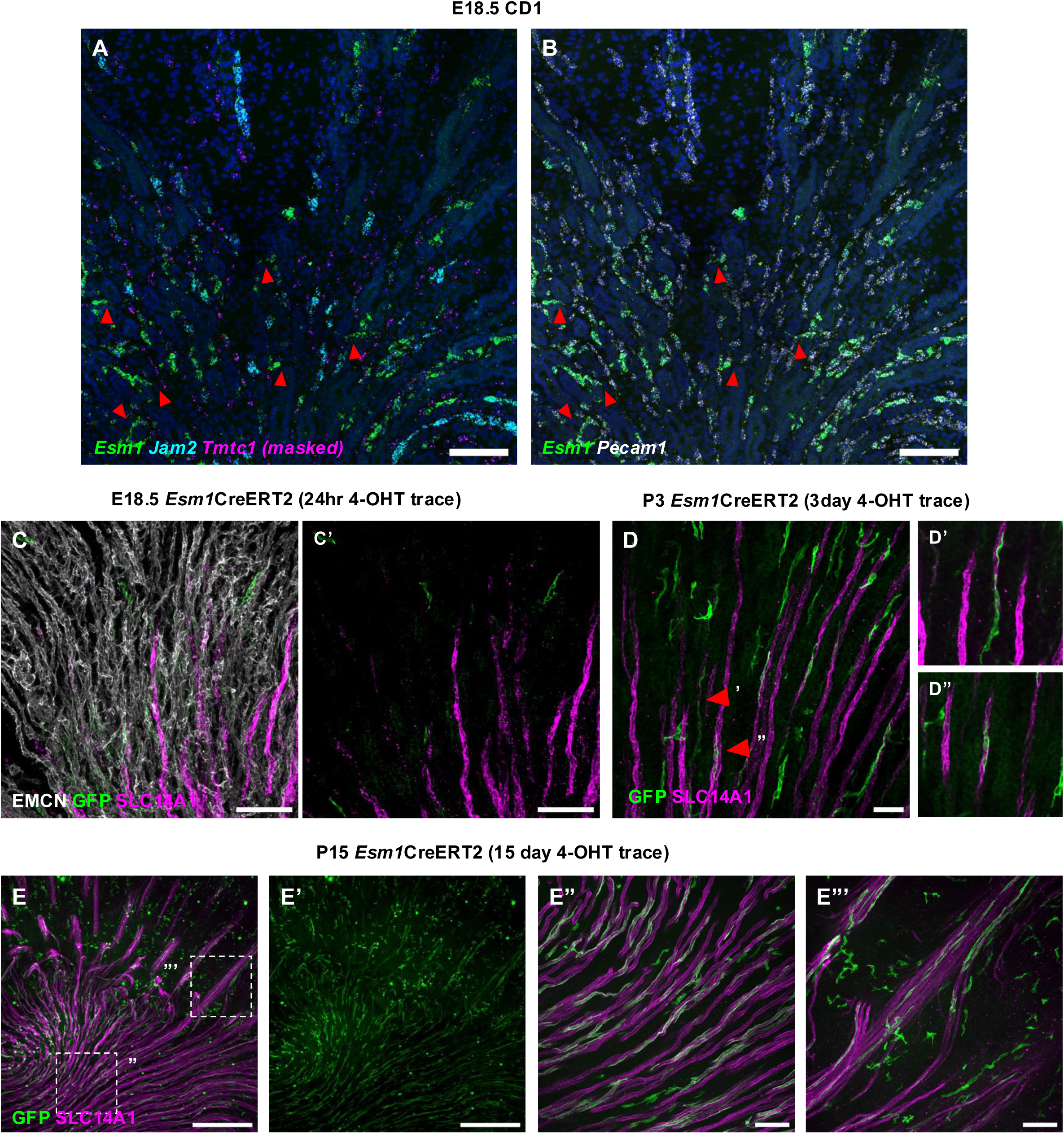
*Esm1* lineage cells in the medulla contribute to the descending vasa recta. (A) *Esm1*, *Jam2,* and *Tmtc* (masked with *Pecam1*) RNAscope on medullary section of E18.5 kidney. (B) *Esm1* and *Pecam1* RNAscope on section from (A). (C) WMIF of medulla of E18.5 *Esm1*CreERT2; mT/mG^f/f^ 24 hour lineage trace kidney for EMCN, GFP, and SLC14A1. (C’) GFP and SLC14A1 only view of (C). (D) WMIF of medulla of P3 *Esm1*CreERT2; mT/mG^f/f^ 3 day lineage trace kidney for GFP and SLC14A1. (D’-D”) Zoom views of (D) showing individual vessels. (E) WMIF of medulla of P15 *Esm1*CreERT2; mT/mG^f/f^ 15 day lineage trace kidney for GFP and SLC14A1. (E’) GFP only view of (E). (E”) Zoom view of (E), showing inner medulla. (E”’) Zoom view of (E), showing outer medulla. Scale bars: 100µm (A-C, E”-E”’), 50µm (D), 500µm (E).

**Data S1.**
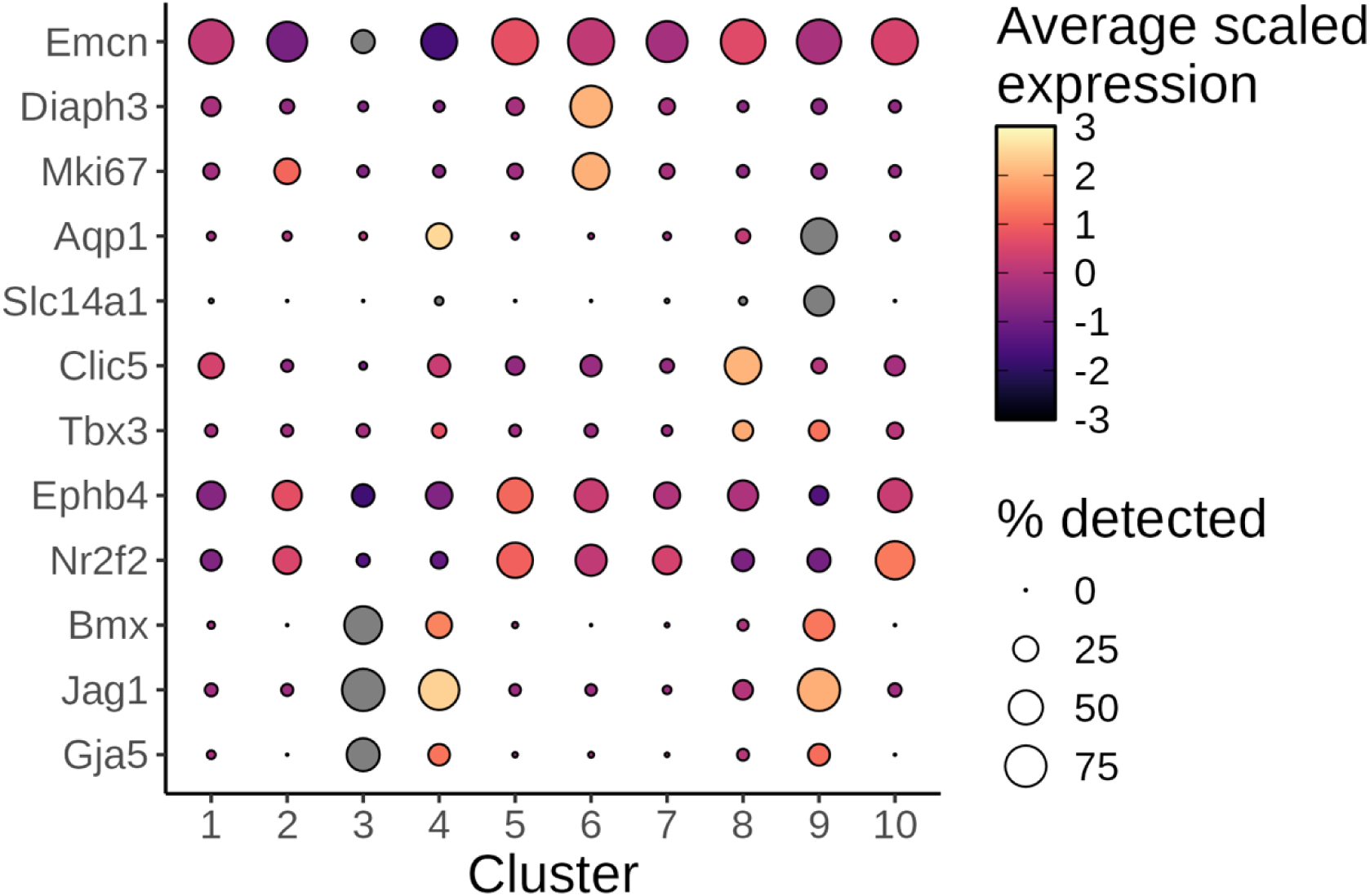
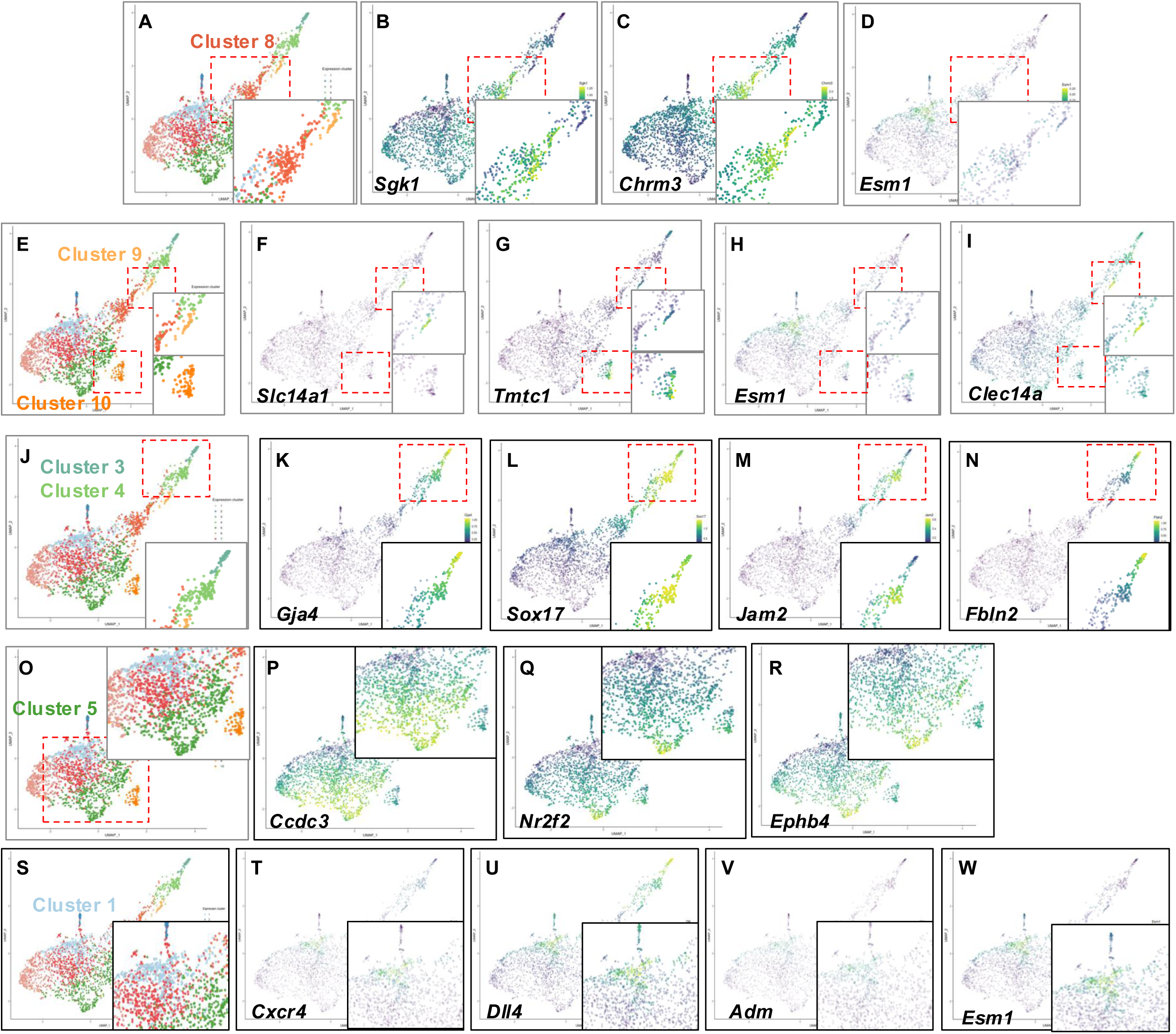
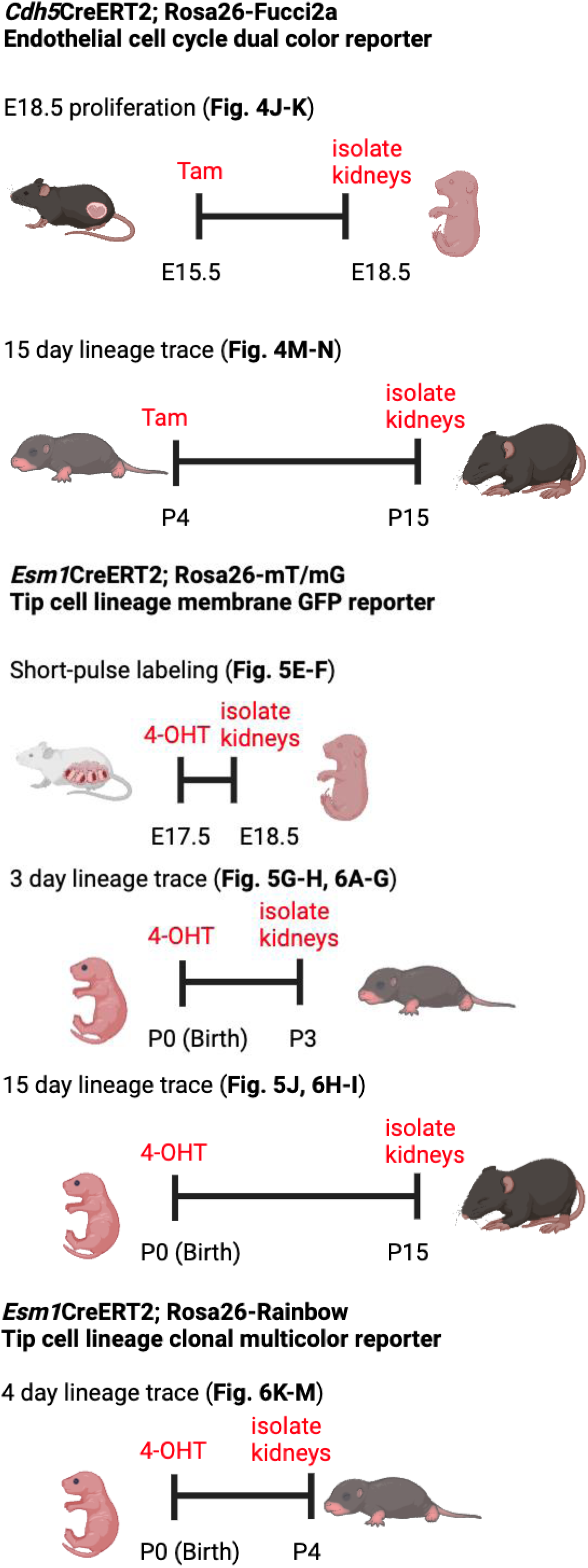
Additional bioinformatic analysis and experimental information, related to Figures 1, 2, 4, 5, 6. 1. Dot plot with additional marker genes, associated with Figure 1 2. UMAP projections of endothelial gene expression, associated with Figure 2, 4, and 5. 3. Schemes for mouse experiments, associated with Figure 4, 5, and 6.

## Notes

### Competing Interest Statement

The authors have declared no competing interest.

